# Fasting primes small intestinal regeneration after damage via a microbiome–metabolite-chromatin axis

**DOI:** 10.64898/2026.03.06.710208

**Authors:** Praveen Barrodia, Ajay Kumar Saw, Sabrina L. Jeter-Jones, Chia-Chi Chang, Jiansu Shao, Emre Arslan, Anand K Singh, Suresh Satpati, Robert R. Jenq, Kunal Rai, Helen Piwnica-Worms

## Abstract

Fasting enhances small intestinal regeneration after radiation but the contribution of the gut microbiome to this process remains uncharacterized. We identify *Akkermansia muciniphila* (*AKK*) as a key mediator of this response. *AKK* was enriched in fasted mice and its antibiotic depletion abrogated radioprotection whereas reintroduction restored both organismal survival and intestinal integrity. Fasting elevated propionic acid, consistent with *AKK*’s metabolic output. *AKK*-conditioned medium and propionate induced histone H3 acetylation in intestinal stem cell cultures while in vivo fasting induced *AKK*-dependent H3K27ac and H3K9ac, remodeling promoter-enhancer landscapes in crypt epithelial cells. Epigenetic profiling revealed a rewired core regulatory program enriched for pioneer transcription factors (Foxa, Gata, Klf), architectural organizers (Ctcf, Boris), and lineage-defining and metabolic regulators (Cdx2, Hnf4). This program supports expansion of a population of persister stem cells characterized by open chromatin accessibility at key stem and regenerative-associated loci including *Clu*, *Olfm4*, *Lgr5, Ascl2, Lrig1, Sox9, Rnf43, and Axin2.* These findings define a fasting-induced microbiome-metabolite-chromatin axis that epigenetically primes highly plastic persister stem cells for rapid regeneration of the intestinal epithelium following radiation-induced injury.

**Significance Statement:** Fasting changes the gut microbiome, but how these changes help the body recover from damage is not well understood. We found that fasting increases a helpful bacterium, *Akkermansia muciniphila*, which produces propionate, which drives epigenetic changes by modifying histones and regulating gene activity. These changes promote the expansion of persister stem cells that help the intestine recover after radiation. This study shows how fasting and gut bacteria work together to protect healthy tissue and suggests that diet or microbial treatments could help reduce side effects of cancer radiotherapy.

## Introduction

Fasting induces widespread changes in metabolism, chromatin architecture and gene expression across multiple tissues^1, 2^ and reshapes the gut microbiota^3^. We previously demonstrated that fasting protects mice from small intestinal injury caused by high-dose etoposide^4^ and ionizing radiation^5^. Fasted mice maintain superior intestinal architecture compared with fed mice shortly after chemotherapy or irradiation, with deeper crypts, taller villi, and reduced epithelial cell damage ^4, 5^, supporting the conclusion that fasting improves tissue structure early after injury. The protection arises in part, by preservation of stem cells in the small intestine (SI), which promotes tissue homeostasis and organismal survival (4,5). However, the molecular mechanisms by which fasting primes the small intestine for regeneration after injury remain poorly understood.

The continuous turnover of the intestinal epithelium is driven by multipotent LGR5⁺ crypt-base columnar cells (CBCs) located at the base of intestinal crypts^6^. However, CBCs are highly sensitive to injury and are rapidly lost following insults such as irradiation^7^. Despite this loss, the intestinal epithelium exhibits a remarkable capacity for regeneration^8^, suggesting the involvement of additional stem or progenitor populations.

A second population of quiescent “+4” cells, also referred to as reserve stem cells (RSCs), was previously proposed to mediate intestinal regeneration following damage^9–12^. Although CBCs and RSCs were initially considered mutually exclusive populations^12, 13^, subsequent studies demonstrated that LGR5⁺ CBCs express markers associated with RSCs^14^. Moreover, functional studies revealed that RSCs are dispensable for intestinal repair, whereas LGR5⁺ cells are essential for regeneration following damage^8^.

More recently, a rare damage-induced stem cell population termed revival stem cells (revSCs) has been identified as a key contributor to intestinal regeneration. revSCs are characterized by high CLU expression and are extremely rare under homeostatic conditions^15^. These cells are multipotent and can give rise to all major intestinal epithelial lineages, including LGR5⁺ CBCs. Following irradiation-induced injury, revSCs undergo a transient YAP1-dependent expansion, reconstitute the LGR5⁺ CBC compartment, and are required for restoration of a functional intestinal epithelium.

In addition to revSCs, multiple progenitor populations—including absorptive enterocyte progenitors^16^, secretory lineage progenitors^17–21^, and slow-cycling LGR5⁺ cells^22^, have been shown to contribute to epithelial regeneration. The transcriptional regulator YAP1, a critical mediator of intestinal regeneration, has been proposed to promote a pro-survival phenotype in LGR5⁺ cells^23^. Collectively, these findings raise an unresolved question: whether intestinal regeneration is driven primarily by specialized injury-responsive stem cell populations or by broader cellular plasticity across multiple epithelial lineages and whether the regenerative response of the small intestine can be preconditioned by stress-associated states such as fasting.

*AKK*, a mucin-degrading commensal bacterium, is enriched in the intestinal microbiome of fasted mice^24^. Here we show that *AKK* is required for fasting-mediated SI radioprotection. Depletion of *AKK* abrogated radioprotection, whereas reintroduction of *AKK* restored epithelial integrity and improved survival. Mechanistically, *AKK* produces the short chain fatty acid propionate, which promotes histone H3 acetylation at lysine 9 and 27 in SI crypt epithelial cells, thereby remodeling enhancer–promoter landscapes. Using integrated single-cell ATAC-seq and H3K27ac CUT&Tag profiling, we identify an *AKK*-dependent transcription factor (TF) regulatory program enriched in Hnf, Foxa, Gata and Klf activities that likely drives expansion of a Clu+Olfm4+ population which we name “persister cells”. This cellular state persists across injury-regeneration phases and is associated with transcriptional programs that accelerate crypt regeneration. Our work provides evidence for revitalization of epigenetic memory of developmental chromatin regulatory programs by regenerative cells post radiation damage and identifies a fasting–*AKK*–metabolite–chromatin axis that primes the intestinal epithelium for injury resilience. Together, these findings establish a microbiome-based framework for protecting normal tissues during radiation therapy.

## Results

### Fasting induces accumulation of *AKK* in the small Intestines of mice

Given that fasting is known to impact the gut flora^24^, we compared the diversity and composition of bacteria in the small intestines (SI) of fed and fasted animals **(Fig.1)**. Mice were allowed to feed freely or were fasted for 24h according to the schema shown in **(Fig. 1a)**. Mice received total abdominal irradiation (TA-XRT) as previously described^25^. Consistent with prior findings^5^, animals fed ad libitum before high-dose radiation died between 7 and 9 days after treatment, whereas all fasted mice survived to the study endpoint **(Fig. 1b)**.

**Fig. 1.**
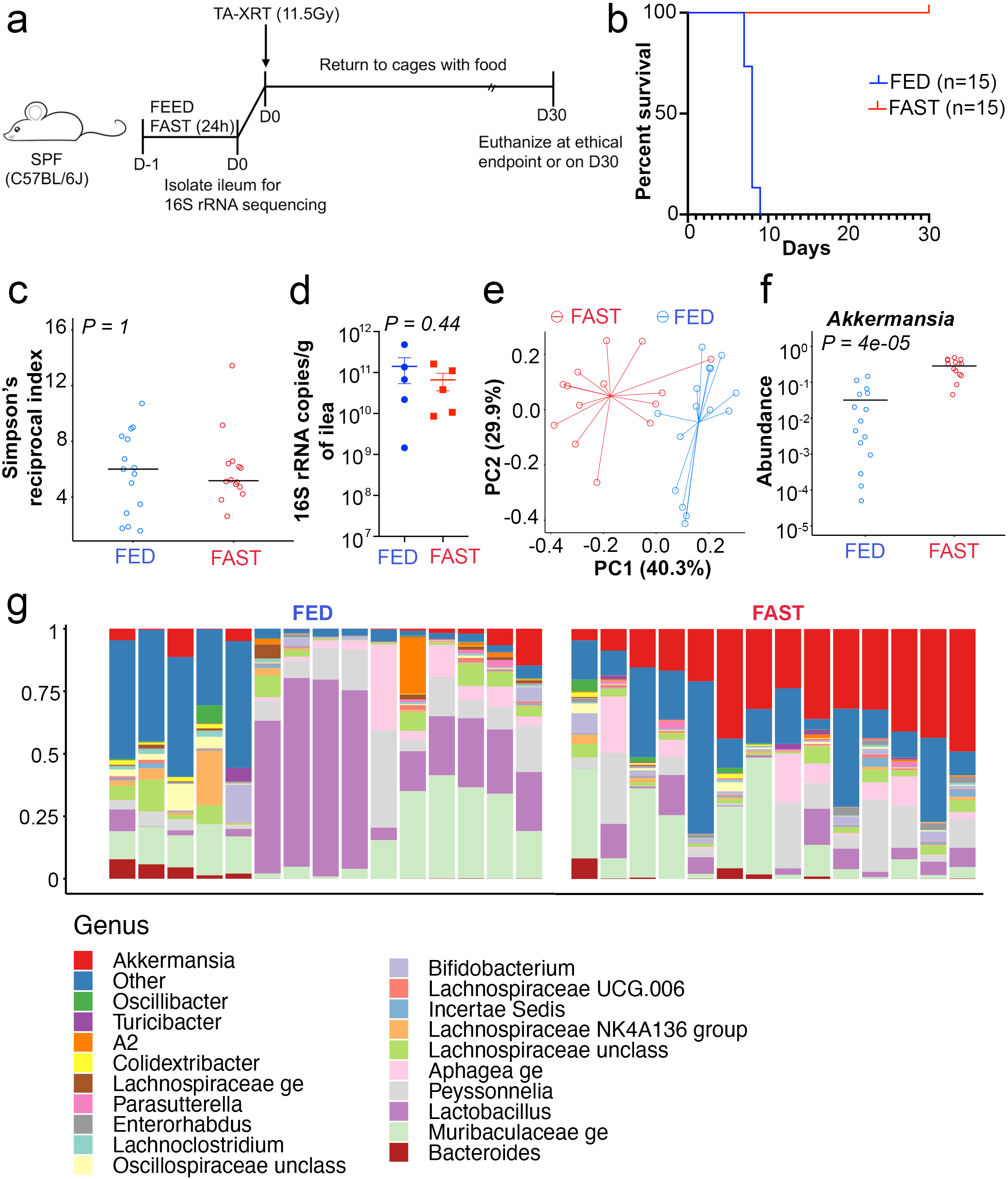
*AKK* accumulates in the small intestines of fasted mice. a. Schematic of the experimental design. C57BL/6J mice were allowed to feed freely or were fasted for 24 h. On day 0, small intestinal ileum samples were collected from one cohort of mice for 16S rRNA sequencing to analyze the SI microbiome composition. Another cohort of mice was exposed to total abdominal radiation (TA-XRT) at a dose of 11.5 Gy. After radiation exposure, mice were returned to single-housed cages with food and survival was monitored daily. b. Survival of fed and fasted mice following TA-XRT was monitored daily for 30 days. P= <0.0001 (Logrank (Mantel-Cox) test) c. Diversity of the SI microbiome in ileum samples collected on D0 was quantified using Simpson’s reciprocal index. *P=1* (using Mann-Whitney U test) d. Absolute abundance of total 16s copies per gram of ileum sample collected from fed and fasted mice were measured by standard spike-in method. Representative data shown from 5 mice. P=0.44. Data are presented as mean ± SEM, analyzed by Student’s t test, unpaired. e. Principal Coordinates Analysis (PCoA) based on weighted UniFrac distances was performed to evaluate the differences in microbiome composition between fed and fasted mice. Statistical significance was determined by permutational MANOVA testing. *P=0.001* f. Relative abundance of *AKK* in ileum samples of fed and fasted mice. *P = 4e-05* (Mann-Whitney U test) g. Relative abundance of bacteria at the genus level in ileum samples collected on day 0. Each column represents an individual mouse.

Assessment of microbiome using 16S rRNA based gene sequencing of fecal material collected from the ilea of fed and fasted mice revealed minimal changes in the diversity **(Fig. 1c)** or absolute abundance **(Fig. 1d)** of bacterial flora between the two groups. However, significant compositional differences between fed and fasted cohorts were readily detected **(Fig. 1e),** with *AKK* abundance significantly enriched in the small intestines of fasted mice compared with their fed counterparts **(Fig.1f, g)**. Other changes in bacterial flora associated with fasting included a significant increase in the relative abundance of *Erysipelatoclostridium*, *Clostridium sensu stricto 1*, *Coprococcus*, and *Marvinbryantia* and a significant decrease in the relative abundance of *Lachnospiraceae*, *Lactobacillus*, *Streptococcu*s, and the *Clostridia vadin BB60 group* **(Supplementary Fig. 1)**.

### *AKK* contributes to organismal and small intestinal radioprotection

To determine if *AKK* contributes to organismal and SI radioprotection, mice were exposed (or not) to broad spectrum antibiotics (Ampicillin and Enrofloxacin) in their drinking water for 7 days according to the schema shown in **(Fig. 2a)**^26^. Microbiome analysis revealed significant reductions in both the diversity **(Supplementary Fig. 2a)** and composition **(Supplementary Fig. 2b, c)** of bacteria in the small intestines of antibiotic treated mice. Antibiotics were allowed to clear from the remaining cohorts of mice and either *AKK* or vehicle (PBS) were introduced by gavage. 16s rRNA sequencing of Ilea fecal material demonstrated that *AKK* successfully colonized the gut of antibiotic treated mice **(Fig. 2b)**. Other cohorts of mice were then mock-irradiated or exposed to TA-XRT at 11.5 Gy and SIs were isolated 5 days later for histology, while remaining mice were followed for 30 days to monitor organismal survival.

**Fig. 2.**
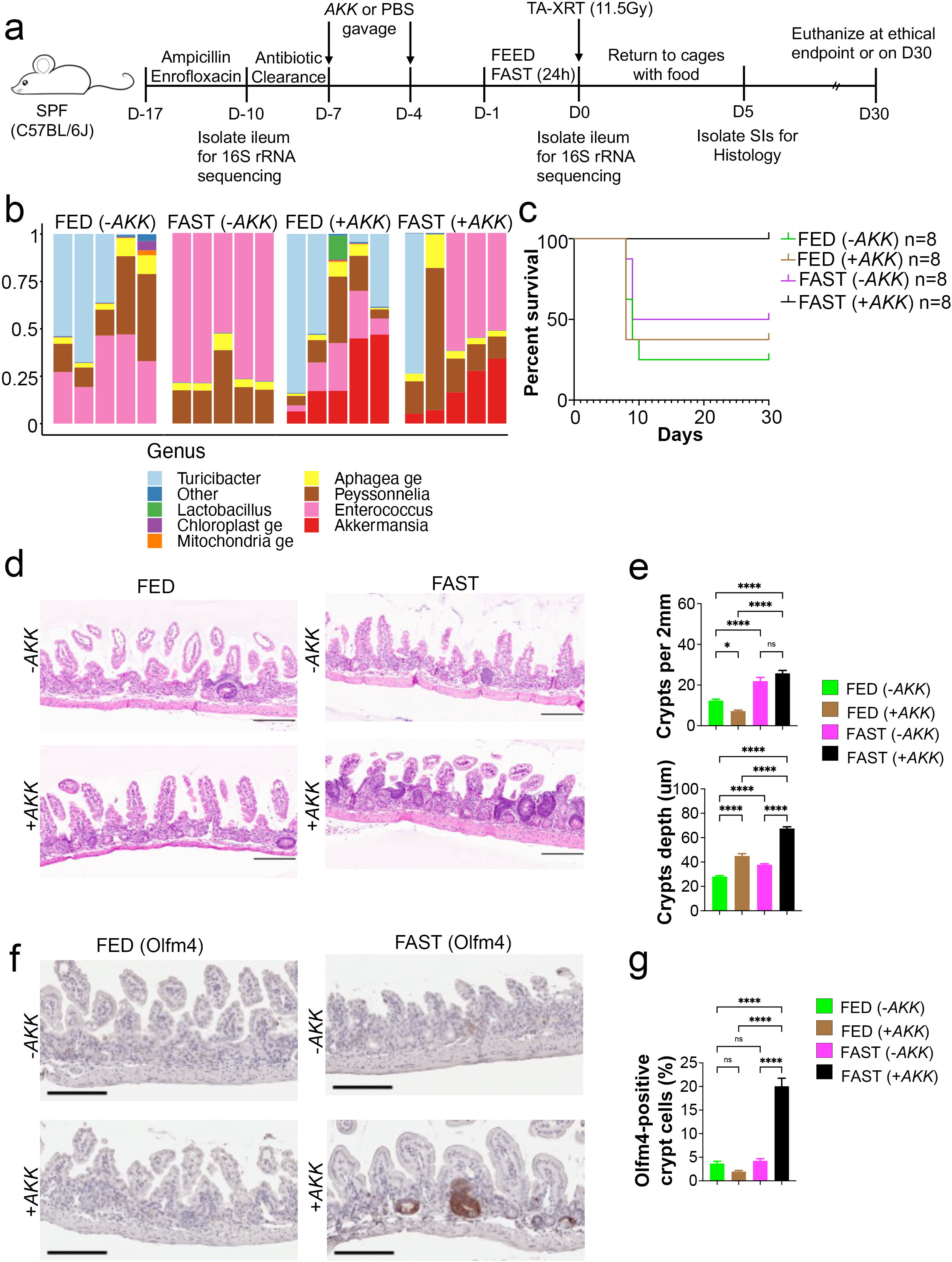
Fasting and *AKK* are both required to protect mice and preserve small intestinal architecture from high dose irradiation. a. Schema of experimental design. C57BL/6J mice were administered antibiotics (Ampicillin 0.5 g/L and Enrofloxacin 0.25 g/L) in drinking water for 7 days followed by three days of antibiotic clearance. *AKK* (1×10^8^ CFU) or vehicle (PBS) were introduced by gavage on day -7 and day -4. On day -1, mice were fed or fasted for 24 h and then exposed to total abdominal radiation (11.5 Gy; D0). Mice were then housed singly with food and on day 5 post radiation a cohort of mice was sacrificed, and ileums isolated for analysis. Remaining mice were monitored for survival until day 30 post radiation. Ileums were also isolated from mice on D-10 and D0 for 16S rRNA sequencing. b. Relative abundances of bacteria at the genus level in ileum samples collected on D0. Each column represents an individual mouse. c. Survival of mice following TA-XRT was monitored daily for 30 days. P= 0.0023 for FED (-*AKK*) vs FAST (+*AKK*), P= 0.8488 for FED (-*AKK*) vs FED (+*AKK*), P= 0.2698 for FED (-*AKK*) vs FAST (-*AKK*) (using Logrank (Mantel-Cox) test) d. Representative images of H & E-stained ileum collected on Day 5. Scale bars, 200 µm. Magnification, 20x. Representative images shown from 4 mice. e. Number of crypts per 2 mm of SI (ileum) and their depth were quantitated for each day 5 sample. The average number of crypts per mm of ileum length and the average crypt depth were plotted (n=10 fields per mouse). ns nonsignificant; *P<0.05; ****P<0.0001 by one-way ANOVA. Error bars are ± SEM. f. Representative images of ileum stained for Olfm4 (day 5 samples). Scale bars, 200 µm. Magnification, 20x. Representative images shown for 4 mice. g. Percentage of crypt epithelial cells staining positive for Olfm4 is shown. ns, nonsignificant; ****P<0.0001 by one-way ANOVA. Error bars are ± SEM. 10 fields per mouse form 4 mice.

Disruption of the SI microbiota through antibiotic treatment impaired the ability of fasting to protect mice from high dose radiation as only 50% of antibiotic treated fasted mice survived to study endpoint **(Fig. 2c)**. Importantly, reconstitution with *AKK* fully restored the ability of fasting to provide radioprotection to antibiotic treated mice as 100% of these mice survived to study endpoint. Although *AKK* levels varied among fasted mice **(Fig. 2b)**, all fasted animals survived irradiation **(Fig. 2c)** suggesting that a threshold level of colonization is sufficient to confer radioprotection. Interestingly, reconstitution with *AKK* did not significantly improve the survival of antibiotic treated fed mice **(Fig. 2c)**. Normally 100% of fed mice die within 6 to 10 days after exposure to high dose radiation. Interestingly, this was not observed in fed mice that received broad-spectrum antibiotics prior to radiation exposure, suggesting that antibiotics provide partial protection from the loss of the barrier function which is compromised by high-dose radiation in fed mice. Importantly, the SIs of fasted antibiotic-treated mice reconstituted with *AKK* exhibited a significant increase in crypt depth **(Fig. 2d, e)** as well as significantly more Olfm4+ stem cells **(Fig. 2f, g)** relative to antibiotic treated fasted cohorts not reconstituted with *AKK.* Conversely, these increases in crypt depth and Olfm4+ stem cells were not observed in fed antibiotic-treated mice reconstituted with *AKK*. Taken together, these results suggest that *AKK* contributes to fasting-induced radioprotection at both the organismal and SI levels.

### Specific depletion of *AKK* from the gut microbiota disrupts fasting-induced radioprotection

Whereas broad spectrum antibiotics significantly disrupted the gut microbiome **(Fig.2)**, a three-week course of tetracycline has been shown to deplete *AKK* without specific depletion of other bacterial species^27, 28^. Therefore, we treated mice with tetracycline to determine if loss of *AKK* followed by its reconstitution impacted host responses to high dose radiation **(Fig. 3a)**. A three-week course of tetracycline exposure in drinking water was sufficient to deplete *AKK* from the gut microbiome of mice **(Supplementary Fig. 3a, b)**. 16S rRNA gene sequencing revealed changes in the composition of bacteria in the small intestines of tetracycline treated mice **(Supplementary Fig. 3c)**, but not to the same extent as observed in mice treated with broad spectrum antibiotics **(Supplementary Fig. 2c)**. Importantly, tetracycline pre-treatment impaired the ability of fasting to radioprotect and reconstitution with *AKK* was sufficient to fully restore radioprotection to fasted animals **(Fig. 3b, c)**. By contrast, reconstitution with *AKK* did not improve survival of similarly treated fed animals. Analysis of small intestinal integrity revealed a significant increase in crypt number, crypt depth and Olfm4+ stem cells in fasted mice reconstituted with *AKK* compared with all other groups **(Fig. 3d-g)**. Taken together, these data suggest an essential role for *AKK* in fasting-mediated radioprotection.

**Fig. 3.**
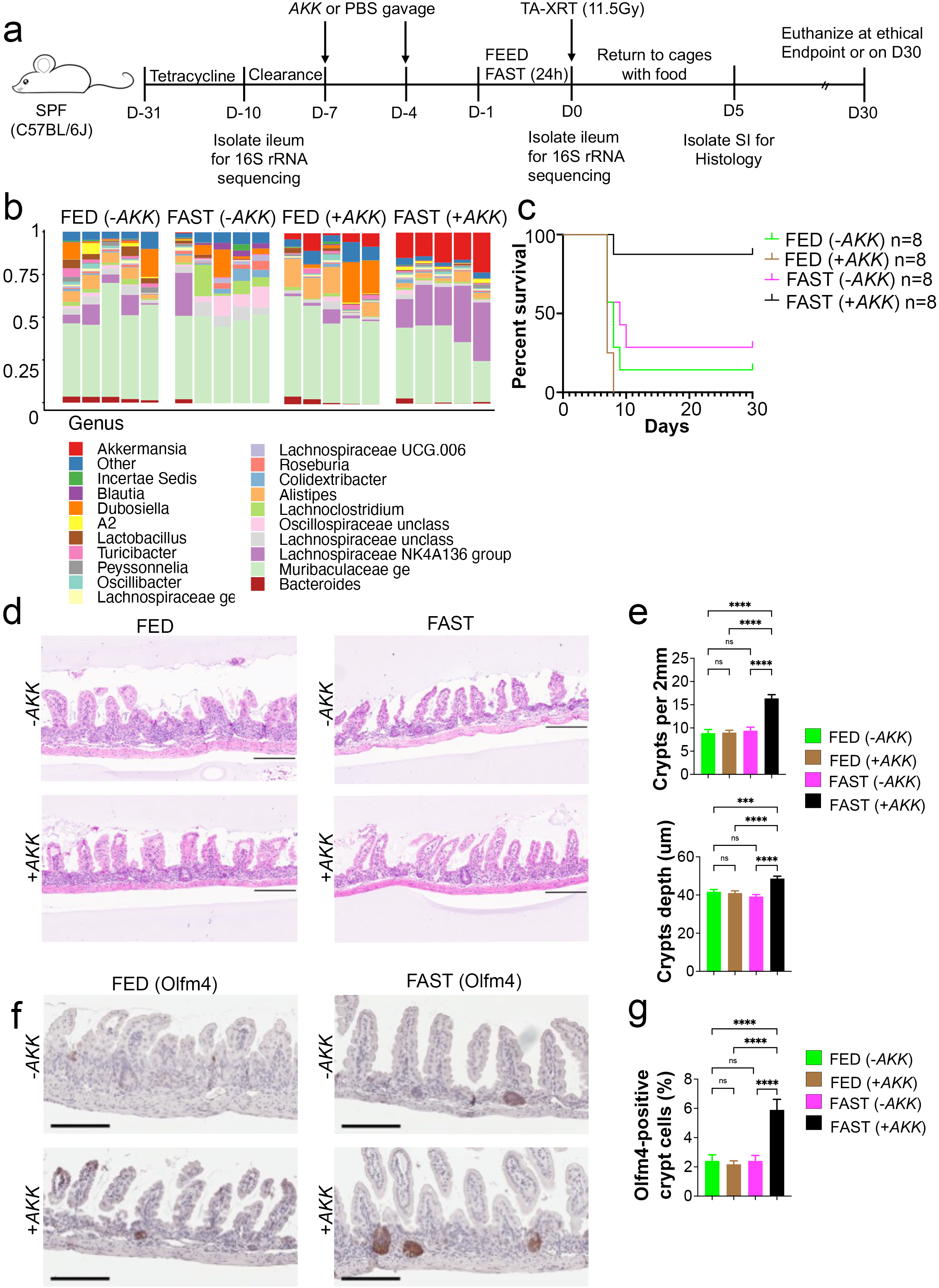
Specific elimination of *AKK* hampers fasting mediated radiation protection in mice. a. Schema of experimental design. C57BL/6J mice were administered vehicle or antibiotics (Tetracycline 3 g/L and 10% sucrose) in drinking water for 21 days followed by three days of antibiotic clearance. *AKK* (1×10^8^ CFU) or vehicle (PBS) were introduced by gavage on day -7 and day -4. On day -1, mice were fed or fasted for 24 h and then exposed to total abdominal radiation (11.5 Gy; D0). Mice were then housed singly with food and on day 5 a cohort of mice was sacrificed, and ileums isolated for analysis. Remaining mice were monitored for survival until day 30 post radiation. Ileums were also isolated from mice on D-10 and D0 for 16S rRNA sequencing. b. Relative abundances of bacteria at the genus level in ileum samples collected at D0. Each column represents an individual mouse. c. Survival of mice following TA-XRT was monitored daily for 30 days. P= 0.0038 for FED (-*AKK*) vs FAST (+*AKK*), P= 0.1036 for FED (-*AKK*) vs FED (+*AKK*), P= 0.4313 for FED (-*AKK*) vs FAST (-*AKK*) (using Logrank (Mantel-Cox) test) d. Representative images of H & E-stained ileum collected on Day 5. Scale bars, 200 µm. Magnification, 20x. Representative images shown from 4 mice. e. Number of crypts per 2 mm of SI (ileum) and their depth was quantitated for each day 5 sample. The average number of crypts per mm of ileum length and the average crypt depth was plotted (n=10 fields per mouse). ns, nonsignificant; ***P<0.0003; ****P<0.0001 by one-way ANOVA. Error bars are ± SEM. f. Representative images of ileum stained for Olfm4 (day 5 samples). Scale bars, 200 µm. Magnification, 20x. Representative images shown for 4 mice. g. Percentage of crypt epithelial cells staining positive for Olfm4 is shown. ns, nonsignificant, ****P<0.0001 by one-way ANOVA. Error bars are ± SEM. 10 fields per mouse form 4 mice.

### *AKK* metabolite promotes histone H3 acetylation in small intestinal epithelial cells

*AKK* is a gram-negative anaerobic bacterium that degrades and utilizes mucus produced by intestinal goblet cells^29^. Single cell RNA sequencing demonstrated that goblet cells became significantly enriched in the small intestines of fasted animals relative to their fed cohorts **(Fig. 4a-c, Supplementary Fig. 4)** and this likely accounts for the enrichment of *AKK* in the gut microbiome of fasted animals. The mucin degradation activity of *AKK* leads to the production of short chain fatty acids (SCFAs) including propionate and acetate, which can be used by some non-mucus-degrading bacteria to produce butyrate^30^.

**Fig. 4.**
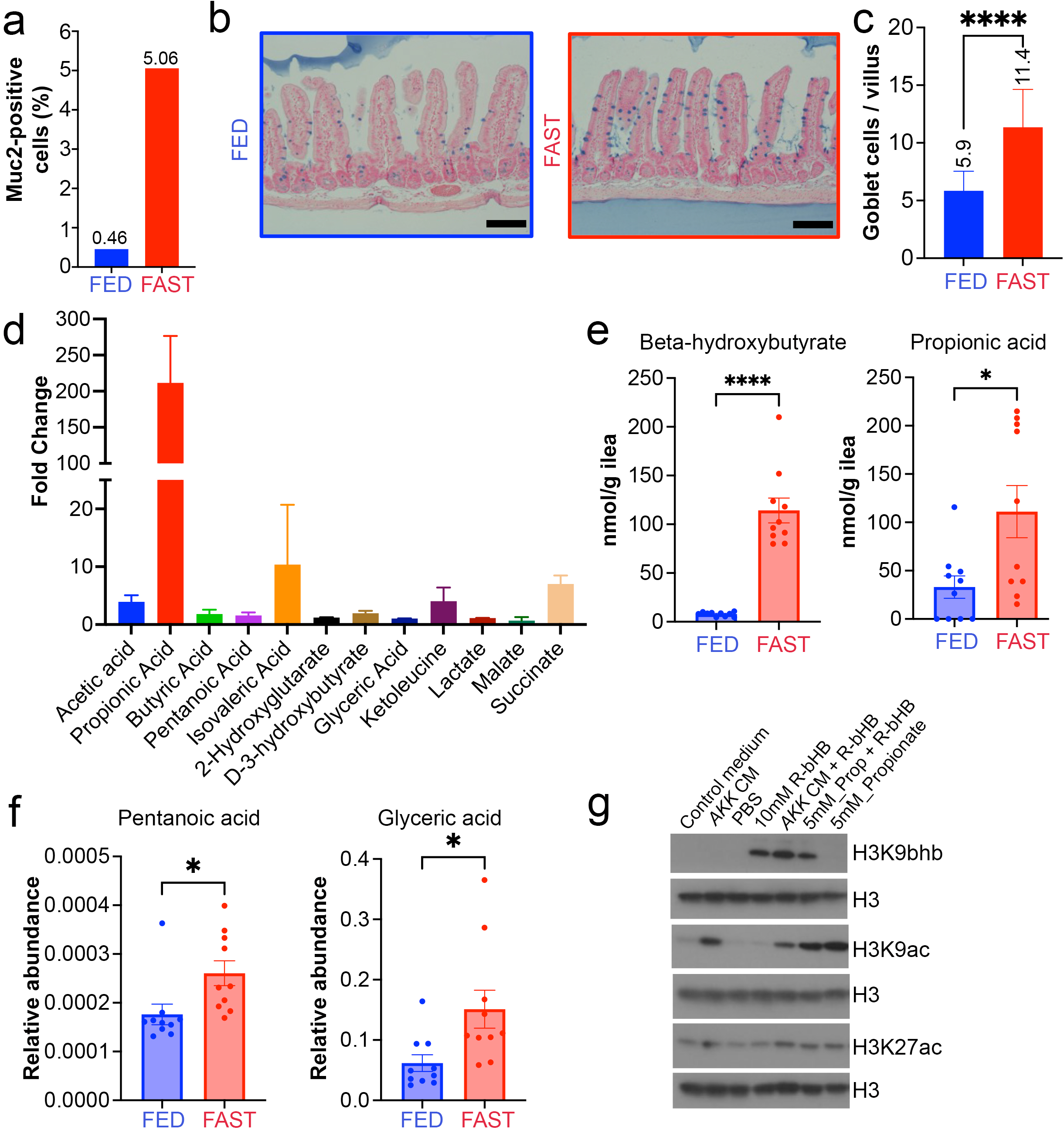
Fasting and *AKK* induces epigenetic changes in SI epithelial cells. a. C57BL/6J mice were allowed to feed freely or were fasted for 24 h. Small intestinal cells were isolated and subjected to single cell RNA-Sequencing. Percentage of Goblet cells, identified by *Muc2* expression, was quantified. Total 4034 cells form fasted and 3724 cells from fed group. b. Ileum samples from fed and 24 h fasted mice were stained with Alcian Blue to identify Goblet cells. Scale bars, 100 µm. Magnification, 20x. Representative images shown from n=4 mice. c. The number of Goblet cells in Ileum samples from fed and fasted mice was determined by counting Alcian Blue positive cells in 10 fields from 4 mice. Data are presented as mean ± SEM, analyzed by Student’s t test, unpaired. d. Murine *AKK* (MDAJAX AM001) was cultured in brain heart infusion media containing 5 mg/ml of mucin under strict anaerobic conditions. Relative abundance of short-chain fatty acids measured in *AKK* conditioned- and control-media was determined by mass spectrometry and is presented as relative fold change. e. Ileum samples were isolated from fed (n=10) and 24h fasted (n=10) mice and absolute concentrations of b-hydroxybutyrate and propionic acid were determined using Ultra-High-Resolution IC-MS. Data are presented as mean ± SEM, analyzed by Student’s t test, unpaired. *P=0.0161 and ****P=<0.0001. f. Relative abundance of ileum SCFA concentrations in fed (n=10) and 24h fasted mice (n=10). Data are presented as mean ± SEM, analyzed by Student’s t test, unpaired. *P<0.02 g. Representative western blot analysis of H3K9bhb, H3K9ac, and H3K27ac of acid-extracted histones from stem cell-enriched epithelial spheroids generated from small intestinal crypts. Spheroids were treated with (R)-(–)-3-hydroxybutyric acid (R-bHB) for 24 h, or for 3 h with other listed components. 2 mg of histones were loaded per well, and histone H3 was used as the loading control.

Metabolic profiling identified propionic acid as the most significantly enriched metabolite in *AKK-* conditioned media compared with control media **(Fig. 4d)**. In addition, metabolic profiling of ileal contents from fed and fasted mice revealed elevated levels of propionic acid and beta-hydroxybutyrate (β-OHB) in fasted animals relative to fed controls **(Fig. 4e)**^31^. In addition, pentanoic acid, a microbial fermentation product related to propionic acid, and glyceric acid, an intermediate in carbohydrate and glycerol metabolism, were significantly increased in the ilea of fasted animals **(Fig. 4f)**. The accumulation of these microbial- and host derived- metabolites points to a coordinated fasting-induced reprogramming of gut microbial fermentation and intestinal energy metabolism.

SCFAs regulate diverse cellular functions including fueling SI goblet cells to enhance mucin production and thereby promote *AKK* accumulation. They also modulate epigenetic and transcriptional programs either directly, through histone acetylation or propionylation^32^, or by serving as substrates for butyrate-producing bacteria to inhibit histone deacetylases (HDACs)^33^. β-OHB also functions as a chromatin regulator both via lysine β-hydroxybutyrylation (Kbhb) of histones H3 and H4^31, 34^ and by inhibiting class I HDACs^35^. Consistent with this, *AKK-*conditioned medium and its major metabolite propionate induced histone H3 acetylation in stem cell-enriched SI epithelial spheroids cultured in vitro **(Fig. 4g)**. Moreover, Kbhb enrichment was observed in these cultures following treatment with the bioactive β-OHB derivative R-bHB.

### *AKK* induces epigenetic changes in small intestinal epithelial cells, contributing to protection against radiation

We previously reported that fasting alters specific chromatin states of SI epithelial cells in vivo leading to upregulation of specific pathways involved in adaptation to nutrient deprivation^31^. To dissect the contribution of *AKK-*derived metabolites to this process, we mapped active enhancers and promoters by CUT&Tag profiling of the histone acetylation marks H3K27ac and H3K9ac in SI crypt epithelial cells under fasting and *AKK-*depleted conditions. Mice were divided into fed (Fed) and fasted (Fast) groups and each group was treated with or without tetracycline (-/+ Tetra) to selectively deplete *AKK* **(Fig. 5a)**. Fasted mice harbored significantly increased average intensity and number of H3K27ac and H3K9ac peaks (Fast -Tetra vs Fed-Tetra). **(Fig. 5b-c, Supplementary Fig. 5a, b)**. This effect was significantly reduced when *AKK* was depleted by tetracycline in the fasted group (Fast +Tetra), However, *AKK* depletion showed minimal effect on acetylation peak enrichment in fed mice (Fed + Tetra). Overlap analysis identified 37,601 H3K27ac peaks and 88,040 H3K9ac peaks (p value =10^-5^) that were uniquely present under the Fast (–Tetra) condition, indicating an *AKK* dependent effect on many regulatory elements. The most pronounced changes were observed in H3K27ac, followed by H3K9ac **(Fig. 5c, Supplementary Fig. 5b)**. Together, these data suggest a key role for *AKK* in regulating fasting-induced histone acetylation patterns in SI crypt epithelial cells.

**Fig. 5.**
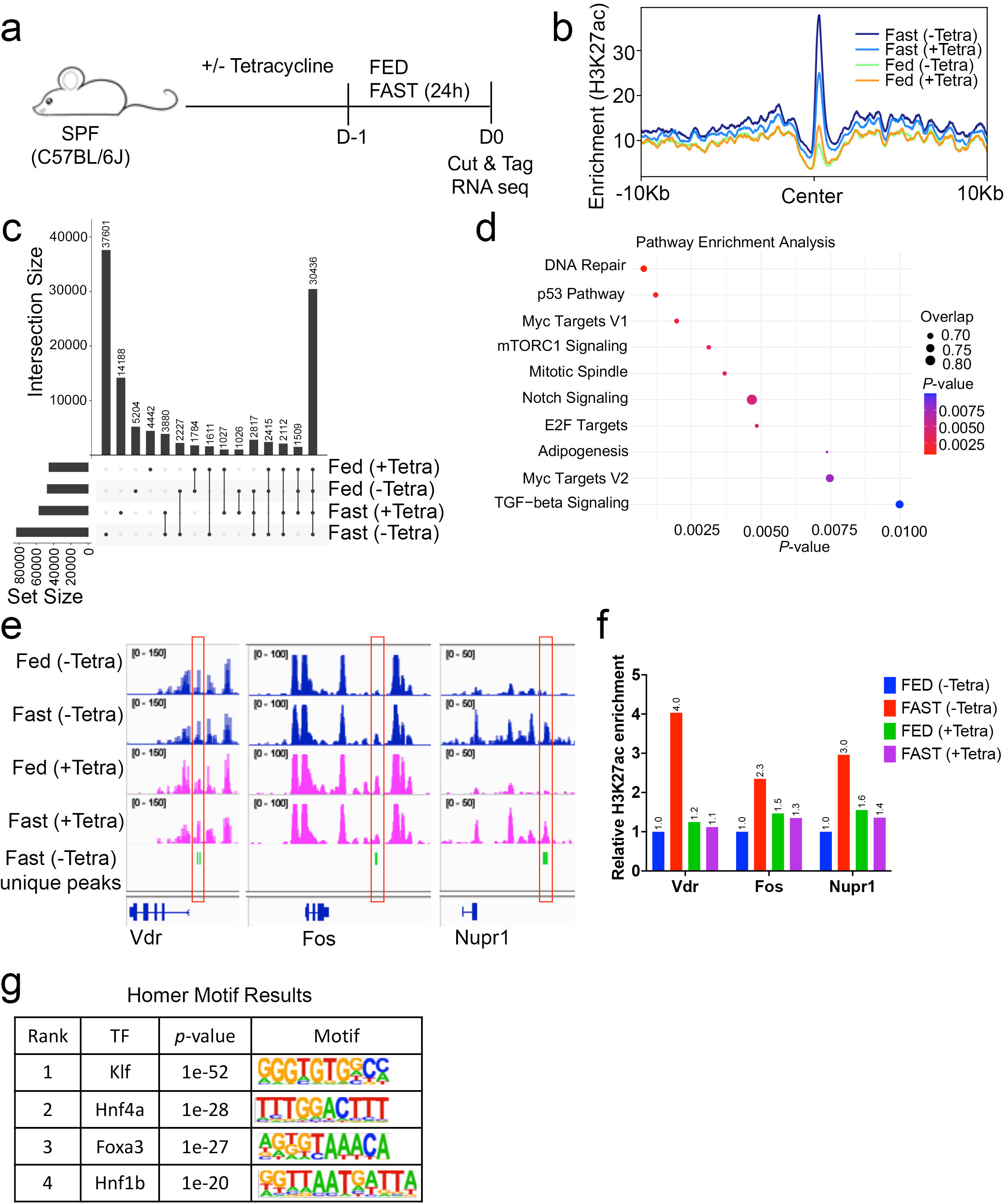
*AKK* differentially enriches H3K27ac at proximal promoters of apoptotic and proliferative program in small intestine crypts. a. Schematic of whole-crypt isolation for CUT&Tag and bulk RNA seq analyses. b. Enrichment plot for H3K27ac peaks in Fed (-Tetra), Fed(+Tetra), Fast (-Tetra) and Fast (+Tetra). c. Upset plots showing overlap of CUT&Tag seq H3K27ac peaks among FED (-Tetra), Fed (+Tetra), Fast (-Tetra) and Fast (+Tetra). The CUT&Tag seq peaks unique to Fast (-Tetra) were analyzed separately. d. Top 10 MSigDB GSEA pathways based on H3K27ac proximal promoter peaks overlapping with publicily available HiChIP and inhouse ChIP seq data in Fast (-Tetra) condition. e. Genome browser view of CUT&Tag tracks for H3K27ac-enriched regions in SI crypts at the p53 pathway-responsive genes Vdr, Fos and Nupr1 under FAST (-Tetra) conditions. f. ChIP-qPCR analysis of H3K27ac enrichment at *Vdr, Fos*, and *Nupr1*-specific peaks in Fed (-Tetra), Fed (+Tetra), Fast (-Tetra) and Fast (+Tetra) samples. g. Motif enrichment analysis for enhancers unique to Fast (-Tetra) group using HOMER.

For gene annotation, we generated enhancer–promoter loop interaction data by integrating publicly available HiChIP datasets^36^ with in-house H3K27ac CUT&Tag and ChIP-seq data^31^ collected under both Fed and Fast conditions. Functional enrichment analysis of uniquely present H3K27ac and H3K9ac peaks in the Fast (-Tetra) group showed significant associations with pathways related to DNA repair and cell proliferation. Notably, genes involved in the p53 signaling pathway were highly represented **(Fig. 5d, Supplementary Fig. 5c)**. These included well-characterized p53 target genes such as *Vdr*, *Fos*, and *Nupr1*, which showed increased H3K27ac and H3K9ac signals in the Fast (-Tetra) group compared to both Fed groups and the *AKK* depleted Fast (+Tetra) group **(Fig. 5e, Supplementary Fig. 5d)**. H3K27ac ChIP–qPCR confirmed increased H3K27ac enrichment at each p53 target gene in the fasted group compared to all other groups, including the FAST (+ Tetra) group, supporting a requirement for *AKK* in the establishment of this chromatin mark under fasted conditions **(Fig. 5f)**.

To better understand the core regulatory programs that drive the radiation protection, we analyzed H3K27ac-marked enhancer regions unique to the Fast (-Tetra) group using motif enrichment analysis. This revealed a significant enrichment of binding sites for the KLF (Klf), HNF (Hnf4a, Hnf1b), and FOXA (Foxa3) transcription factor (TF) families **(Fig. 5g)**. Klf and Foxa3 are well-characterized pioneer factors capable of opening compacted chromatin and establishing enhancer activity during gut tube development and specification^37^.

### Accumulation of Clu+ Olfm4+ persister cells in SI crypts of fasted animals

To investigate how fasting shapes the chromatin landscape and regenerative potential of the intestinal epithelium following injury, we performed single-cell ATAC-seq (scATAC-seq) on crypt epithelial cells isolated from fed or 24 hour fasted mice exposed to TA-XRT (11.5 Gy). Samples were collected at baseline (0 h) and at 24-, 48-, or 72-hpi **(Fig. 6a)**. UMAP projections of aggregated scATAC-seq data revealed distinct clustering by sample **(Fig. 6b, Supplementary Fig. 6a)** and by cell type **(Fig. 6c, Supplementary Fig. 6b)**, with fasting associated with pronounced shifts in chromatin accessibility. We identified two clusters (3 and 6) characterized by accessibility at the *Clusterin* (*Clu*) locus. *Clu* expression characterizes revival stem cells important for intestinal regeneration following irradiation^15^. Marker gene accessibility analysis distinguished these two clusters. Cluster 3 displayed high *Clu* accessibility with no accessibility at *Olfm4,* whereas Cluster 6 showed accessibility at both the *Clu* and *Olfm4* loci along with stronger accessibility at the *Ascl2* and *Hmgcs2* loci (**Fig. 6d**). Notably, in addition to *Clu* and *Olfm4*, Cluster 6 exhibited accessibility at multiple intestinal stem cell marker genes, including *Lgr5*, *HopX*, *Lrig1*, *Sox9* **(Fig. 6e)** and IPA canonical pathway enrichment analysis identified Wnt/β-catenin signaling and stem cell pluripotency-associated pathways as significantly upregulated in C6 cells relative to C3 cells (**Supplementary Fig.6c).** Analysis of genes associated with open chromatin regions in C6 identified *Sox21, Nr5a2, Hnf1a, Fzd2, Axin2, and Ccnd1* revealing strong enrichment of Wnt, Notch, and other pathways critical for intestinal regeneration **(Supplementary Fig.6d)**. Analysis of genes associated with open chromatin regions in C3 identified *Tcf4, Arrb2*, *Ago1*, *Hao1*, and *Spr*, indicating enrichment of metabolic pathways in addition to modest enrichment of Wnt pathway **(Supplementary Fig.6d).**

**Fig. 6.**
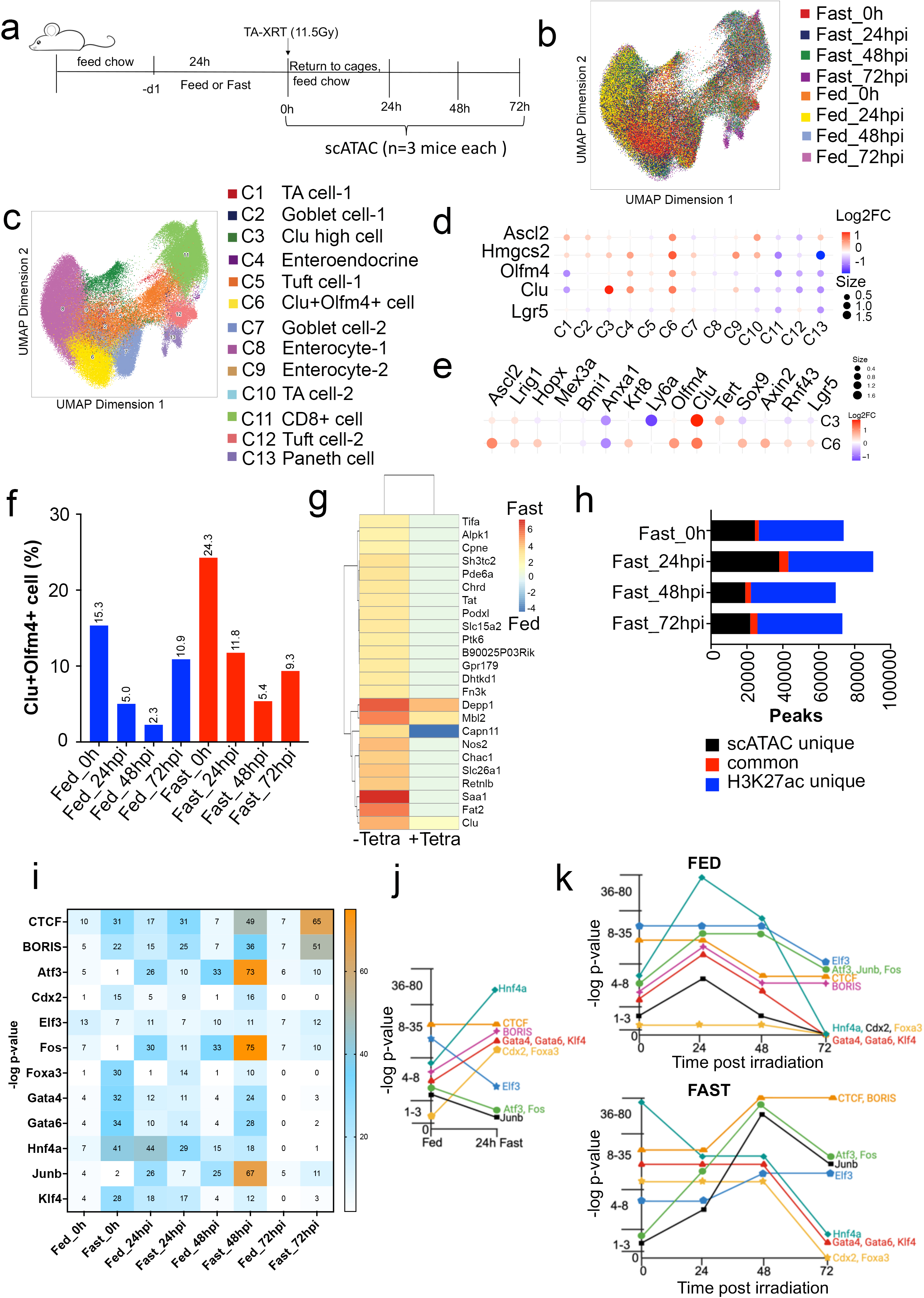
Fasting remodels the chromatin landscape of intestinal stem and revival-like cell populations. a. Experimental design. Mice were fed ad libitum or fasted for 24 h, followed by total abdominal irradiation (11.5 Gy). Small intestinal crypt cells were collected at 0 h, 24 hpi, 48 hpi, or 72 hpi (post-irradiation) and subjected to single-cell ATAC sequencing (scATAC-seq). b. UMAP projection of aggregated scATAC-seq data colored by sample, showing condition-specific shifts in chromatin accessibility. c. UMAP colored by cluster identity, identifying major intestinal epithelial populations. d. Bubble plot showing chromatin accessibility of stem and revival marker loci (Clu, Olfm4) across conditions. Cluster 6 corresponds to the Clu+Olfm4+ persister stem cell population. e. Bubble plot showing chromatin accessibility (log2FC) of stem cell, revival, and injury-associated gene loci in Cluster 3 (C3) and Cluster 6 (C6). f. Percentage of Clu+Olfm4+ cells (Cluster 6) across all samples, demonstrating increased representation during fasting. g. Integration of CUT&Tag H3K27ac data with RNA-seq for the activated genes in Fed (-Tetra), Fed (+Tetra), Fast (-Tetra) and Fast (+Tetra) samples. All genes displayed are differentially expressed (p ≤ 0.05, log2FC ≥ 2 or ≤ –2). n = 2 mice per group. h. Bar graph showing the number of shared accessible regions between scATAC-seq and H3K27ac CUT&Tag datasets. “Common” refers to overlapping enhancer-like peaks. i. HOMER motif enrichment analysis showing -log10 p-values for transcription factor motifs associated with intestinal progenitor identity and injury response (IR) protection, alongside their corresponding gene expression levels from bulk RNA-seq in fed and fasted conditions and at 0-, 24-, 48-, and 72-hours post irradiation (hpi). j. Graphical representation of data shown in panel i for fed and 24 fasted mice. Y axis represents binned -log10 p-values of transcription factor motif activity. k. Graphical representation of data show in panel i for fed and fasted mice. mice at 0-, 24-, 48-, and 72-hours post irradiation (hpi). Y axis represents binned -log10 p-values of transcription factor motif activity.

Clusters 3 and 6 also differed in their temporal dynamics following irradiation (**Fig. 6f** and **Supplementary Fig.6e**). Cluster 6 was enriched in mice fasted for 24h at baseline and more abundant following irradiation at 24 and 48 hpi before converging toward similar proportions by 72 hpi. In contrast, Cluster 3 represented a smaller and relatively stable population over time with only modest changes at the 24 and 48 hpi time points. By 72 h cluster 3 declined to 1.2% in fasted mice but remained at 3.1% in fed mice. Collectively, these data demonstrate that Clusters 3 and 6 represent biologically distinct epithelial cell populations with divergent chromatin states and regenerative dynamics with preferentially expansion of Cluster 6 in response to fasting. ScRNA sequencing supported the conclusion that fasting enriches for stem cell and revival stem cell markers **(Supplementary Fig. 6f)**.

Next, to assess fasting and *AKK* associated transcriptional changes in SI crypts, we identified differentially expressed genes (DEGs; p ≤ 0.05 and a log_2_ fold change cutoff of ≥ 2 or ≤ –2) by bulk RNA-Seq. Analysis was conducted on fed and 24h fasted mice treated or not with tetracycline. DEGs exhibiting enrichment of H3K27ac or H3K9ac peaks within 5 kb upstream or 1 kb downstream of their transcription start sites were significantly upregulated under fasting conditions **(Fig. 6g, Supplementary Fig. 6g)**. Notably, *Clu* was selectively expressed in the crypts of fasted animals, and its expression was markedly reduced in tetracycline-treated fasted animals indicating that *AKK* contributes to *Clu* induction during fasting.

Trajectory (**Supplementary Fig. 7**) and marker gene accessibility (**Fig. 6d, e**) analyses distinguished cluster 3 from canonical revival stem cells. revSCs arise in the intestine only following irradiation and are characterized by expression of *Clu* and *Ly6a* ^15, 38^. In contrast, cluster 3 cells are present prior to irradiation and do not exhibit significant Ly6a chromatin accessibility. Cluster 3 may represent a transitional revSC-like state, rather than a fully defined revSc population. Fasting primarily expanded the Clu+Olfm4+ population (**Fig. 6f**) which is consistent with an injury responsive cell pool that supports regeneration.

To gain insight into the regulatory programs that drive persistence of the Clu+Olfm4+ persister cell population (C6) upon radiation, we integrated scATAC-seq data with H3K27ac CUT&Tag profiles, which identified 4,042, 3,093, 5,157, and 2,149 “common” open active enhancer peaks in 0 h, 24 hpi, 48 hpi, and 72 hpi fasted samples, compared to 1,241, 2,100, 1,268, and 421 “common” peaks in the corresponding fed samples **(Fig. 6h, Supplementary Fig. 8a)**.

To identify functional regulators, we filtered enriched transcription factor (TF) motifs based on known roles in gut progenitor identity, injury response (IR) protection, and expression in bulk RNA-seq and highlighted candidate TFs potentially driving the fasting-induced state (**Fig. 6i).** This analysis revealed a marked reprogramming of regulatory networks following 24 hours of fasting **(Fig. 6j).** Fasting strongly increased chromatin accessibility surrounding motifs for transcription factors associated with epithelial cell differentiation and metabolic regulation, most notably Hnf4a, which shifted to the highest activity bin, Motifs for Gata actors (Gata4/6), Klf4, and the intestinal lineage regulators Cdx2 and Foxa3 were also enriched. In addition, motifs for the architectural proteins Ctcf and Boris increased, suggesting fasting-induced remodeling of chromatin organization. In contrast, accessibility of motifs associated with Atf3, Fos and JunB declined to the lowest bin in fasted mice, likely due to nutrient deprivation.

Comparison of TF motif accessibility dynamics following irradiation revealed fundamental differences between fed and fasted states **(Fig. 6k).** Fasted animals displayed specific reprogramming of regulatory programs with persistent accessibility of Gata factors, Klf4, Cfdx2. Foxa3, Hnf4a up to 48 hpi and sustained Ctcf/Boris accessibility through 72 hpi. Notably, motifs associated with Atf3, Fos and JunB declined to the lowest accessibility bin after 24 h of fasting **(Fig. 6i)** and robustly rebound when animals were returned to food immediately after irradiation (0 hpi) and resumed feeding **(Fig.6k).** A pronounced increase in AP-1 family motif chromatin accessibility was observed in irradiated fasted animals **(Fig. 6k**) which likely reflects a combined effect of refeeding and radiation-induced signaling. In contrast, in fed animals, motif accessibility either persisted (Elf3, Ctcf) or transiently increased by 24 hpi, but most motifs lost accessibility by 48 hpi, and all motifs lost accessibility by 72 hpi, potentially due to excessive cell death.

Single-cell correlation data showed that cells with high *Clu* accessibility also exhibited elevated chromatin accessibility at regulatory regions associated with Olfm4, Gata4, Gata6, Sp1, Klf4, Foxa3, and Elf4, transcription factors **(Supplementary Fig. 8b).** These results indicate that the fasting-induced transcriptional reprogramming likely occurs in Clu+Olfm4 persister stem cells.

Collectively, these data indicate that fasting reprograms transcriptional regulatory programs critical for intestinal stem cell persistence, metabolic adaptation and regenerative competence following irradiation.

## Discussion

Here we report that *AKK* becomes enriched in the microbiome of fasted mice leading to chromatin remodeling in small intestinal crypt cells. This occurs in a programmatic fashion leading to the reprogramming of key regulatory TF networks to establish a transcriptional output in persister stem cells that promote regeneration of the small intestine following high dose radiation.

Our functional studies demonstrated that *AKK* recolonization is sufficient to confer radioprotection in antibiotic-treated fasted mice. Notably, the radioprotection conferred by *AKK* was restricted to the fasting context, as reintroduction of *AKK* into antibiotic-treated fed mice did not confer protection. These results demonstrate that *AKK* is necessary but not sufficient to confer radioprotection. Beyond reshaping the microbiome, fasting induces β−OHB which contributes to the histone modifications and associated transcriptional reprogramming in small intestinal crypt cells ^31^. These changes, together with other fasting-induced adaptations, are absent in fed mice and likely explain their lack of radioprotection by the addition of *AKK* alone.

We also found that fasting enriched members of the Furmicutes including *Erysipelatoclostridium*, *Clostridium sensu stricto 1*, *Coprococcus*, and *Marvinbryantia,* with all but *Erysipelatoclostridium* known to produce butyrate (but not propionate)^39, 40^. Unlike *AKK*, these taxa are not consistently enriched across fasting studies^24^. While it remains possible that these additional taxa contribute complementary metabolic or signaling functions, our data support *AKK* as the principal microbial mediator of the fasting phenotype. Future studies may clarify if these taxa interact with *AKK* or influence epithelial responses to injury. Interestingly, a previous study showed that *Enterococcus* and *Lachnospiraceae can* provide protection against whole-body radiation even in the absence of fasting, with tryptophan identified as a key metabolite mediating this radioprotective effect (41), In our mouse cohort, *Enterococcus* was not detected and although *Lachnospiraceae* was present at low levels, its abundance did not change in response to fasting.

The regenerative response of the small intestine to radiation injury has been defined by the acute loss of Lgr5+ stem cells, followed by activation of reserve, revival stem and progenitor cells that transiently sustain renewal. Prior studies have implicated p53, YAP/TAZ, ASCL2 activities and TGFβ, Notch and Hedgehog pathways in this revival function^15, 38, 41^. These canonical pathways collectively restore crypt integrity, but they are only engaged after injury.

Our findings indicate that fasting does not replace established regenerative programs but instead primes the intestinal epithelium before injury through epigenetic reprogramming, thereby enhancing the regenerative response once DNA damage occurs. This fasting-induced epigenetic priming does not alter intestinal architecture on its own. In our prior studies villus height, crypt depth, and crypt numbers were shown to be indistinguishable in the SIs of fed and fasted mice^4, 5^. Architectural differences emerged only after irradiation, with fasting leading to better preservation of crypt structure, consistent with radioprotection.

Fasting increased goblet cell numbers which supplies mucin, thereby enriching *AKK* in the gut microbiota. *AKK* produces propionate, while host metabolism contributes β-hydroxybutyrate (β-OHB). Together, these metabolites function as chromatin cofactors, inducing H3K27ac, H3K9ac, and β-hydroxybutyrylation^31^, thereby remodeling enhancer–promoter interactions. This remodeling selectively increases accessibility at loci bound by pioneer transcription factors including Foxa, Gata and Klf families^42, 43^. Foxa and Gata are classical pioneer factors that bind condensed chromatin, displace nucleosomes, and establish enhancer competence in endodermal development^44^. Their enrichment in SI crypts of fasted animals suggests that fasting re-engages developmental pioneer programs to establish regenerative competence prior to injury.

Clu+ revival cells are a rare, injury induced population that emerge after intestinal injury, marked by *clusterin* (*Clu)* expression, activation of a fetal-like gene program including *Ly6a*, and activation of Yap/Hippo and IL6/STAT3 pathways. They serve as a transient regenerative reserve, replenishing lost Lgr5+ stem cells and contributing to epithelial repair^15, 38^. The fasting-induced Clu+ cell populations identified in our study (Clusters 3 and 6) are distinct from canonical Clu+revSCs. Unlike revSCs, which emerge only after injury and express *Ly6a,* the Clu-high (Cluster 3) and Olfm4+Clu+ (Cluster 6) cells we describe are present in fasted animals prior to irradiation and exhibit low (Cluster 3) or absent (Cluster 6) *Ly6a* chromatin accessibility.

Based on maker gene accessibility-, and trajectory-analyses, Clu-high, (Cluster 3) cells may represent a transitional, revSC-like population. This population is enriched by fasting but does not expand during the injury responsive period. Clu+Olfm4+ (Cluster 6) cells are also enriched by fasting, but they expand during the early injury-response period and exhibit enrichment of several SI stem cell markers. These properties are consistent with an injury responsive persister stem cell pool that supports regeneration.

Epigenetic profiling revealed a rewired core regulatory program in fasted mice that primes injury resilience, characterized by pioneer transcription factors (Foxa/Gata/Klf) that pre-emptively license chromatin accessibility; lineage-defining and metabolic regulators (Cdx2, Hnf4) that coordinate metabolic programs required for stem cell survival and regenerative capacity and architectural organizers (Ctcf, Boris) that stabilize higher-order chromatin structure.

Our data support a hierarchical model in which pioneer TFs first establish chromatin accessibility, enabling a subsequent binding of settler TFs (Cdx2 and Hnf2) at their cognate regulatory elements. Both the enhancer and promoter of Clu harbor conserved Hnf and Cdx binding motifs, directly linking fasting-induced chromatin remodeling to the regulation of Clu+ stem cell states^45^. Architectural organizers (Ctcf, Boris) then stabilize these higher order, fasting-induced chromatin configuration, thereby priming Clu+ persister stem cells for a rapid response to irradiation-induced injury. In parallel, AP1-components c-jun and Fos support proliferative programs during the regenerative phase.

Overall, we propose a model in which fasting-induced nutrient deprivation elicits a stress response in the small intestine, characterized by reduced epithelial cell proliferation and accumulation of host- and microbiome-derived metabolites. Intestinal crypt cells respond to this metabolic environment by remodeling chromatin accessibility to promote cellular plasticity. Under these conditions, C6 cells accumulate and become epigenetically “primed” for multiple differentiation trajectories. We hypothesize that this enhanced plasticity enables rapid and effective regeneration of the intestinal epithelium following irradiation-induced injury **(Fig. 7).**

**Fig. 7.**
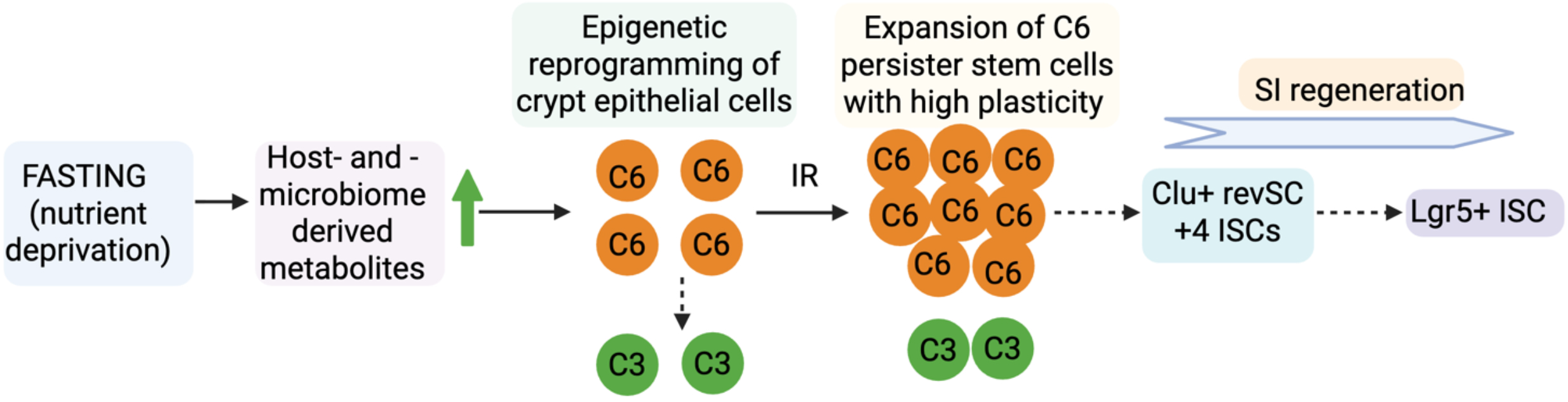
Model summarizing findings.

## Methods

### Study Approval

This study was carried out in accordance with the guidelines outlined in the Guide for the Care and Use of Laboratory Animals from the National Institutes of Health. Ethical considerations regarding animal care and use were approved by the MD Anderson Institutional Animal Care and Use Committee (IACUC) under protocol number 00001101-RN04. The euthanasia of animals followed established criteria set by the Association for Assessment and Accreditation of Laboratory Animal Care International and the IACUC euthanasia endpoints.

### Animals and feeding

C57BL/6J male mice purchased from Jackson Laboratories (JAX, 000664) were maintained at 72°F ± 2°F on a 12-hour light/dark cycle. Male mice aged between four to nine weeks were randomly assigned to either fed or fasted groups. Fed mice had unrestricted access to both food (PicoLab 5053, #0007688) and water for 24 hours, while fasted mice were deprived of food for 24 hours but had unrestricted access to water. Mice were singly housed on aspen bedding for the duration of the experiments. The number of animals used is provided in figure legends.

### Total abdominal radiation

Mice were subjected to a single dose of 11.5 Gray (Gy) of total abdominal radiation therapy (TA-XRT). The radiation treatment was carried out using the X-RAD 225 Cx irradiator, a device designed for delivering controlled and targeted radiation therapy.

### 16S rRNA sequencing

Small intestinal ilea samples were collected from mice and weighed prior to DNA extraction. Synthetic spike-in was done as described^46^. Briefly, *Escherichia coli* DH5α cells (Thermo Fisher, Cat. EC0112) were prepared in various concentrations (10^6^–10^10^ cells) for DNA isolation. Cell counts and viability (98%) were assessed using Syto BC dye and propidium iodide, with automated counting. A plasmid mixture of SP1-pUC57-Kan, SP2-pUC57-Kan, SP3-pUC57-Kan, and SP4-pUC57-Kan was spiked in equal molar ratios (0.5 ng) with 600 μl of InhibitEx lysis buffer (Qiagen, Cat. 19593) and added to bacterial or fecal samples for DNA extraction, followed by bead-beating with metal (Qiagen, Cat. 6999) and zirconia beads (BioSpec, Cat. 11079101Z). Genomic DNA was isolated using the QIAamp Fast DNA Stool Kit (Qiagen, Cat. 51604).

The V4 region of the 16S rRNA gene was amplified by PCR from each extracted and purified genomic DNA using the 515 forward (5’-GTGYCAGCMGCCGCGGTAA) and 806 reverse (5’- GGACTACNVGGGTWTCTAAT) primer pairs. The amplicon pools were purified with a QIAquick gel extraction kit (Qiagen, Cat. 28704) and sequenced on the Illumina MiSeq sequencer using a 2 x 250 bp paired-end protocol. The experiments assessing the impact of different spike-in standards on sequencing run quality used a 2 x 150 bp paired-end protocol. The reads were merged, dereplicated, and length-filtered utilizing VSEARCH.20 Following denoising and chimera calling using the UNOISE3 commands,21 the unique sequences, or zero-radius OTUs (ZOTUs), were taxonomically classified using Mothur.22 with the SILVA database version 138. Alpha and beta diversity metrics were generated in QIIME 2. PCoA analysis using the weighted UniFrac distance assessed by PERMANOVA testing.

### Culturing *AKK*

*AKK* (MDAJAX AM001) was cultured in brain heart infusion agar containing 5 mg/ml of mucin under strict anaerobic conditions. Single colonies were then isolated and cultured in brain heart infusion broth for 48-72 h at 37 °C, under strict anaerobic conditions. *AKK* was diluted to a final concentration of 1×10^8^ CFU in 0.2ml of anaerobic phosphate-buffered saline (PBS) for all mouse experiments.

### PCR amplification of *AKK* specific gene from fecal material

Fecal material collected from small intestinal ilea was resuspended in 100 µL of nuclease-free water and incubated at 96°C with shaking for 7 minutes to lyse bacterial cells and inactivate nucleases. Samples were then centrifuged at 12000 rpm for 5 minutes at 4°C and the supernatant isolated. Primers specific for genes encoding Amuc1483F (5’ GGCGGAGTCATGGTGTATATC 3’) and Amuc1483R (5’ CAGACCGGAGAGAAAGGAATAAA 3’) were used in a PCR reaction using the Q5^®^ High-Fidelity DNA Polymerase. An amplicon of 459 bp was visualized on a 1.5% agarose gel.

### Broad-spectrum antibiotic treatment

C57BL/6J mice were administered antibiotics (Ampicillin 0.5 g/L and Enrofloxacin 0.25 g/L) in drinking water for 7 days followed by three days of antibiotic clearance. *AKK* (1×10^8^ CFU) or vehicle (PBS) were introduced by gavage. Gavaging was repeated 2 days later.

### Tetracycline treatment

C57BL/6J mice were administered vehicle or antibiotics (Tetracycline 3 g/L and 10% sucrose) in drinking water for 21 days followed by three days of antibiotic clearance. *AKK* (1×10^8^ CFU) or vehicle (PBS) were introduced by oral gavage. Gavaging was repeated 2 days later.

### Analysis of Short Chain Fatty Acids by Ultra-High-Resolution IC-MS

Samples collected from small intestinal ilea or *AKK* conditioned media was obtained as described above. Approximately 50 mgs of ilea or 2ml conditioned media were snap frozen in liquid nitrogen, then homogenized with Precellys Tissue Homogenizer. Metabolites were extracted using ice-cold 0.1% Ammonium Hydroxide in methanol:water = 80:20 (v/v). Samples were centrifuged at 17,000 g for 5 min at 4°C, and supernatants were transferred to clean tubes, followed by evaporation to dryness under nitrogen. Dried extracts were reconstituted in deionized water, and 10 μL were injected for analysis by ion chromatography (IC)-MS using the Thermo Scientific (Dionex ICS-6000+) system which includes a Thermo IonPac AS11 column (4 µm particle size, 250 x 2 mm) with a column compartment kept at 35°C and an autosampler tray chilled to 4°C. The IC mobile phase A (MPA; weak) was water, and the mobile phase B (MPB; strong) was water containing 100 mM KOH. A The mobile phase flow rate was 360 µL/min, and the gradient elution program was: 0-2 min, 1% MPB; 2-25 min, 1-40% MPB; 25-39 min, 40-100% MPB; 39-50 min, 100% MPB; 50-50.5 min, 100-1% MPB. The total run time was 55 min. To assist with desolvation for better sensitivity, methanol was delivered by an external pump and combined with the eluent via a low dead volume mixing tee. Data were acquired using a Thermo Orbitrap IQ-X Tribrid Mass Spectrometer under ESI negative ionization mode. Raw data files were imported into Thermo Trace Finder 5.1 software for final analysis. The relative concentration of each compound was normalized to stool weight per sample.

### Isolation of small intestinal crypts

Isolation of small intestinal crypts was conducted following established protocols as previously published^4^ with minor adjustments. Briefly, mice were euthanized by CO_2_ asphyxiation and the entirety of their small intestines (SI) isolated. Intestines were thoroughly flushed with PBS (Ca2+- and Mg2+-free) containing 2 mM EDTA and 100 nM TSA (Trichostatin A) to remove fecal matter. The mesentery was removed, and the SI sample was longitudinally cut and then transversely cut into four equal pieces. Each piece was placed on ice in PBS (Ca2+- and Mg2+-free) containing 100 nM TSA while the rest of the samples were collected. All SI pieces were incubated in PBS (Ca2+- and Mg2+-free) containing 2 mM EDTA and 100 nM TSA for 10 minutes. The samples were then transferred to Hank’s Balanced Salt Solution (HBSS) (Ca2+- and Mg2+-free) containing 2 mM EDTA and 100 nM TSA.

Crypts were released through a series of vortex washes at 1,600 rpm in HBSS (Ca2+- and Mg2+-free) containing 2 mM EDTA and 100 nM TSA, all performed at 4°C. Supernatants from all vortex washes were filtered through a 70-μm mesh to remove villus material and tissue fragments. These filtered solutions were pooled into 50 ml conical tubes. Isolated crypts were pelleted by centrifugation at 1,000 rpm at 4°C. Purified whole crypts were for single-cell sequencing, CUT&Tag experiments, bulk RNAseq, western blots, qPCR, and establishment of spheroid cultures^47^.

### Generation of stem cell-enriched epithelial spheroid cultures

Purified small intestinal crypts were mixed with 30 μL of Matrigel and plated in a 24-well tissue culture dish following the procedure described^47^. Cultures were grown in 50% L-WRN conditioned media supplemented with 10 μM TGF-β RI Kinase Inhibitor VI (SB431542, Calbiochem) and 10 μM Y-27632 dihydrochloride (Millipore Sigma). The culture medium was refreshed every second day, and the spheroids were subcultured every third day for experimental treatments, spheroids were treated with compounds as described in the figure legends.

### Immunohistochemistry and immunofluorescence

Mice small intestinal samples were harvested as described previously^4^ and ilea sections were used for all analyses. Briefly, small intestines were flushed with PBS, fixed in 10% neutral buffered formalin, paraffin-embedded, and cut transversely for subsequent hematoxylin and eosin (H&E) staining was performed by the RHCL or CROR Histology Core. H & E-stained slides were scanned at MD Anderson RHCL using the Aperio AT2 slide scanner (Leica Biosystems), and crypts were manually counted and statistically analyzed using GraphPad Prism 10.

To perform immunostaining, FFPE tissue slides were baked at 65°C for 1 hour, then dewaxed and rehydrated through graded washes in xylene and ethanol. Epitope retrieval was performed by incubating slides in Reveal Decloaker solution (Biocare Medical # RV1000M) at 96°C for 15 minutes using an EZ-Retriever microwave (BioGenex). After washing with PBS, endogenous peroxidase activity was blocked with Dual Endogenous Enzyme Block (Dako # S2003) for 10 minutes at room temperature. Tissues were blocked with Protein Block (Dako # X0909) and/or normal serum (Vector ImmPRESS # MP-7401), followed by overnight incubation with 1:200 dilution of Olfm4 antibody (CST # 39141S) at 4°C. After PBS washes, ImmPRESS® HRP Goat Anti-Rabbit IgG secondary antibody (Vector laboratories # MP-7451) was used for 30min at room temperature. Next, tissue slides were incubated with horseradish peroxidase (HRP) substrate (Vector ImmPACT # SK-4105) to develop the stain. The slides were then counterstained with Hematoxylin QS (Vector # H-3404), dehydrated through graded ethanol-to-xylene washes, and mounted with permanent mounting medium (VectraMount # H-5000). IHC slides were scanned at MD Anderson RHCL using the Aperio AT2 slide scanner (Leica Biosystems), and Olfm4 positive cells in were counted using QuPath^48^ and statistically analyzed using GraphPad Prism 10.

### Histone Extraction

Histones were extracted and purified following methods outlined in Schechter et al., 2007. Briefly, Small intestinal crypts or stem cell-enriched epithelial spheroids were lysed in 1 mL of buffer containing10 mM Tris-Cl (pH 8), 1 mM KCl, 1.5 mM MgCl_2_, 1 mM DTT (Dithiothreitol), and 1 mM PMSF (Phenylmethylsulfonyl fluoride) supplemented with protease and phosphatase inhibitor cocktails. Lysates were centrifuged at 10,000 g for 10 minutes at 4°C. Pelleted nuclei were resuspended in 400ul of 0.2 M H_2_SO_4_ and incubated for 1 hour at 4°C, followed by centrifugation at 16,000 g for 10 minutes at 4°C. The supernatant containing histones was treated with 33% Trichloroacetic acid (TCA), incubated overnight at 4°C and then centrifuged at 16,000 g for 5 minutes at 4°C. Precipitated histones were washed twice with ice-cold acetone, and the pellet was air-dried, then suspended in 100 μL of ultrapure water, and stored at either -20°C or -80°C.

### Resolution and analysis of histones

Purified histones extracted from small intestinal crypts were separated by SDS-PAGE using 4–20% Criterion™ TGX™ Precast Midi Protein Gel (Bio-Rad) transferred to PVDF membrane and analyzed by western blotting using specific histone antibodies H3K9bhb (PTM BioLabs#1250), H3K27ac (Abcam#4729) and H3K9ac (Abcam#4441) and histone H3 (CST# D1H2) for loading control. Proteins were visualized or signal detected using chemiluminesence (ThermoFisher Scientific # 34076) on a G-Box imager (Syngene, USA).

### RNA isolation and quantification from stem cell enriched spheroid culture

Spheroid cultures were lysed in RNA lysis buffer (PureLink™ RNA Mini Kit, ThermoFisher Scientific # 12183018A) and RNA isolated according to manufacturer’s instructions. Total RNA was treated with DNase followed by column purification (RNA Clean & Concentrator-5, Zymo Research # R1013). Total RNA was quantified using a nanodrop spectrophotometer. The undiluted total RNA was then converted into cDNA using the SuperScript™ III First-Strand Synthesis SuperMix (ThermoFisher Scientific # 18080400). Quantitative PCR (qPCR) was performed on undiluted cDNA in duplicate for each primer and probe set using the TaqMan™ Fast Advanced Master Mix for qPCR (ThermoFisher Scientific # 4444557). qPCR data were normalized to an endogenous control, Gapdh. No-template controls were included for each probe set, and no amplification was observed for any samples. The qPCR analyses were conducted on the Applied Biosystems QuantStudio 6 Pro real-time PCR systems, and all assays were performed in biological triplicate to ensure the reliability of the results.

### Single cell RNA sample preparation

Single-cell suspensions (∼10,000 cells/sample) were processed using the 10x Genomics Chromium Controller with Chromium Chip A, Single Cell V(D)J 5′ Gel Beads v1, and Partitioning Oil to generate GEMs. Reverse transcription was performed (53 °C, 45 min; 85 °C, 5 min), followed by GEM breakage, cleanup, and PCR amplification (13 cycles). Amplified cDNA was purified (SPRIselect, Beckman Coulter) and quantified (Agilent 4200 TapeStation). For 5′ gene expression (5′GEX) library preparation, 50 ng cDNA per sample was fragmented, end-repaired, A-tailed, adaptor-ligated, and indexed (Chromium i7 Sample Index, 10x Genomics) with double-sided SPRIselect cleanup after fragmentation and post-PCR.

All libraries were checked for fragment size distribution (Agilent 4200 TapeStation HS D5000 Assay) and quantified (Qubit dsDNA Quantification Kit, Thermo Fisher Scientific). 5′GEX libraries were pooled and sequenced on a NovaSeq6000 S1 (Illumina) targeting 50,000 read pairs/sample. Sequencing was performed using 10x Genomics-recommended parameters for 5′GEX (Read 1: 26 cycles; Read 2: 91 cycles), with a 100 nt format.

### scRNA-seq analysis

Sequences from the Fed and Fasted siEC samples were processed using Cellranger (v 4.0.0, 10x Genomics). The generated filtered cell UMI count matrix was imported into Seurat (v4.3.0) for analysis in R. Doublets were removed using scDblFinder (v1.13.8) with default parameters. Data were normalized, scaled, and variance stabilized using SCTransform (V2) in Seurat. Differentially expressed genes between cell types were identified using FindMarkers in Seurat (p < 0.05, fold change >3).

### RNA sequencing

Total RNA was isolated from small intestinal crypts with PureLink™ RNA Mini Kit (Thermo Fisher Scientific, # 12183018A), eluted in 50 μl of deoxyribonuclease (DNase)/ ribonuclease (RNase)–free water, and treated with DNase I followed by column purification (RNA Clean & Concentrator-5, Zymo Research # R1013). Integrity of RNA was assessed on a 2100 Bioanalyzer Instrument (Agilent Technologies Inc.). Only samples with RNA integrity number (RIN) factor above 7.5 were used for next-generation sequencing (NGS) library preparation.

### RNA sequencing data analysis

We began by assessing the quality of the RNA sequencing data using FastQC for sequencing quality checks. Transcript abundance was quantified with Salmon, using the GENCODE M21 annotation. The resulting quantification files were imported into R for exploratory data analysis. To streamline this process, we used the R package tximeta to import and summarize transcript-level data at the gene level. Differential expression analysis was then performed using DESeq2 with default parameters.

### CUT&Tag

CUT&Tag assays were conducted using the CUTANA CUT&Tag kit (EpiCypher) according to manufacturer’s instructions. Nuclei were extracted from 100,000 small intestinal crypts using a nuclear extraction buffer containing 20 mM Hepes-KOH (pH 7.9), 10 mM KCl, 0.1% Triton X-100, 20% glycerol, 0.5 mM spermidine, and 1× cOmplete Protease Inhibitor Cocktail, and attached to concanavalin A-conjugated magnetic beads activated with bead activation buffer (20 mM Hepes-KOH (pH 7.9), 10 mM KCl, 1 mM CaCl_2_, and 1 mM MnCl_2_). Beads with attached nuclei were incubated with 0.5 μg of either H3K27ac (Thermo Fisher Scientific, # MA5-23516) or H3K9ac (Thermo Fisher Scientific, # MA5-33384) antibodies in Digitonin150 buffer (20 mM Hepes (pH 7.5), 150 mM NaCl, 0.5 mM spermidine, 0.01% digitonin, and 1× complete Protease Inhibitor Cocktail) supplemented with 2 mM EDTA with gentle shaking overnight at 4°C, followed by incubation with 0.5 μg of secondary antibody in Digitonin150 buffer for 30 minutes at room temperature. After two washes with Digitonin150 buffer, the beads were incubated with 2.5 μl of pAG-Tn5 in Wash300 Buffer (20 mM Hepes (pH 7.5), 300 mM NaCl, 0.5 mM spermidine, and 1× cOmplete Protease Inhibitor Cocktail) supplemented with 0.01% digitonin for 1 hour at room temperature. Beads were then washed with Wash300 buffer containing digitonin, followed by incubation in Wash300 buffer with 10 mM MgCl_2_ for 1 hour at 37°C to perform chromatin tagmentation. The beads were then washed with TAPS buffer (10 mM TAPS (pH 8.5) and 0.2 mM EDTA), and DNA was eluted in 5 μl of SDS release buffer (10 mM TAPS (pH 8.5) and 0.1% SDS) for 1 hour at 55°C and quenched with 15 μl of 0.67% Triton X-100. DNA libraries were amplified for 18 to 21 cycles using 25 μl of NEB Next High-Fidelity 2× PCR Master Mix (NEB, #M0541) and 2.5 μl of unique i5 and i7 index primers for Illumina. Paired-end sequencing was performed on the Illumina NextSeq500 MID 150 system (2 × 75 bp). Antibodies used were against H3K27ac (Thermo Fisher Scientific, #MA5-23516) and H3K9ac (Thermo Fisher Scientific, #MA5-33384).

### CUT&Tag Data processing

The preprocessing of raw reads involved several steps. First, FastQC was utilized to evaluate the quality of the reads. Then, Trim Galore was employed to remove adapters if contamination was detected. The filtered reads were subsequently aligned to the mouse reference genome (mm10) using Bowtie 2. The resulting alignment files were downsampled by condition, sorted, and indexed using SAMTOOLS. Peak calling was performed with the Model-based Analysis of ChIP-Seq (MACS) to identify peaks for the sharp histone marks H3K27ac and H3K9ac. The peak enrichment was calculated over a whole genome “input” background with a p-value 1e-5. Finally, for ChIP-seq track visualization, deepTools was used to generate bigWig files by scaling the BAM files to reads per kilobase per million (RPKM), and these were visualized in the Integrative Genomics Viewer (IGV).

### Identification and enrichment of CUT&Tag peaks

The Model-based Analysis of ChIP-seq (MACS) algorithm was used for peak calling with a p-value threshold of 1e-5 to identify H3K27ac and H3K9ac enrichment over “input” background. Unique peaks between the two sample groups were determined using Bedtools by subtracting the shared peak set from the total number of peaks in each condition, based on MACS-generated data. Heatmaps were created using deepTools’ ComputeMatrix and plotHeatmap commands, utilizing input-subtracted bigWig files.

### DiffBind Analysis

The DiffBind R package was utilized to identify differences between two groups of samples. For each histone modification mark, consensus peaks across all samples were determined using the dba and dba.count functions with default settings. A matrix of normalized read counts across all samples was generated using the dba.normalize function. Differential binding affinity analyses were then conducted using the dba.analyze function.

### Assigning CUT&Tag and ChIP-seq peaks to genes

To assign CUT&Tag and ChIP-seq peaks to genes, we generated an enhancer-promoter loop interaction file using publicly available HiChIP datasets^36^ (SRR11548370, SRR11548371) along with 30 lab-generated CUT&Tag and ChIP-seq samples^31^, which included Fed and Fast conditions. For the HiChIP samples, paired-end reads were aligned to the mm10 genome, duplicates were removed, and reads were assigned to MboI restriction fragments. Valid interactions were filtered, and interaction matrices were created using HiC-Pro. HiC-Pro-filtered reads were further processed, and interaction loops were identified using hichipper. The identified loop interaction files from hichipper were overlapped with MACS peak files of various histone modification marks (H3K27ac, H3K9ac) for both Fed and Fast conditions to construct the enhancer-promoter loop interaction file. For gene annotation, we downloaded the “mm10.refGene.txt.gz” file from the UCSC Genome Browser (http://hgdownload.soe.ucsc.edu/goldenPath/mm10/database/refGene.txt.gz) and generated 5 kb upstream and 1 kb downstream BED files using the bedtools flank command.

### Motif analysis, TF enrichment and pathway analysis

To identify potential transcription factor (TF) motifs enriched in regions of interest, we used the findMotifsGenome.pl command from the HOMER software package. The analysis was performed using a BED file of peak regions as input, with the parameter –size 200 to focus on a 200-bp window surrounding the peak center. To assess the functional significance of the bound peaks, we employed the Genomic Regions Enrichment of Annotations Tool (GREAT) and the Enrichr package, using the mm10 genome as the background and default settings for other parameters.

### Single cell ATAC sample preparation

Nuclei were isolated from ∼1 × 10⁶ cells using a modified 10x Genomics “Nuclei Isolation for ATAC” protocol (CG000169), lysed in chilled lysis buffer, washed, and resuspended in 1× Nuclei Buffer targeting 10,000 nuclei. Quality and concentration were assessed via Trypan blue staining and microscopy. scATAC-seq libraries were prepared using the Chromium Next GEM Single Cell Multiome ATAC + Gene Expression Reagent Kits (CG000338) per manufacturer’s instructions, including nuclei transposition, GEM generation/barcoding, post-GEM cleanup, and sample index PCR, followed by double-sided SPRIselect cleanup. Libraries were quality-checked (Agilent 4200 TapeStation HS D1000) and quantified (Qubit), then pooled and sequenced on a NovaSeq6000 S1 (Illumina) at ∼25,000 read pairs/nuclei using 10x Genomics-recommended parameters (Read N1: 50 cycles; i7 Index: 8 cycles; i5 Index: 24 cycles; Read 2: 91 cycles).

### Single cell ATAC analysis

Single cell multiome fastq files were aligned to the mouse (mm10) genome, cell barcodes were demultiplexed, and UMIs corresponding to genes were counted using the cellranger-arc (v2.0.2) count command using default parameters. Single-cell ATAC-seq data were processed using ArchR (v1.0.2) pipeline^49^ in R (v4.2.2) with the mm10 genome. Fragment files from Cell Ranger ARC outputs were used to generate Arrow files (minTSS = 4, minFrags = 1,000). Predicted doublets were identified (addDoubletScores()) and removed (filterDoublets()). Dimensionality reduction was performed via iterative latent semantic indexing (varFeatures=25,000, dimsToUse=30, sampleCells=5000), followed by Louvain clustering (resolution = 0.2) and UMAP embedding with default parameter. Marker genes and peaks were identified using Wilcoxon rank-sum tests (FDR ≤ 0.01, log2FC ≥ 1.25) and visualized in heatmaps and UMAP projections. Peaks were called on pseudo-bulk replicates grouped by cluster using MACS2 (v2.2.9.1). We used the ArchR pipeline for most of our single-cell ATAC-seq datasets analysis. Integration of scATAC-seq and H3K27ac CUT&Tag peaks was performed using Bedtools^50^ and homer for motif analysis.

### Chromatin immunoprecipitation (ChIP)-qPCR

S mall intestinal crypts were isolated from mice and cross-linked with 1% formaldehyde for 10 min at 37°C, followed by quenching with 125 mM glycine for 5 min at 37°C. Crypts were then collected, lysed, and processed as previously described ^31^. Immunoprecipitated DNA was purified, dissolved in water and analyzed by qPCR using primers targeting the FAST (-Tetra) H3K27ac specific peaks.

Vdr (For) 5’-TGTGGAGAGTCTGCCAGGAT-3’, Vdr (Rev) 5’-TCTTGTTCTTCTGCCCACCC-3’,

Fos (For) 5’-CATGAACCTGTTCGTGCCAG-3’ Fos (Rev) 5’-GCTGTCTCGTTGAACTGTTGT-3’,

Nupr1 (For) 5’-ATGTTGTCCAGGTTGGCCTT-3’ Nupr1 (Rev) 5’-CCTGGCAGAGAGCAAGAGAG-3’,

GeneDesert (For) 5’-ACCAAGCACAGAAAAGGTTCAAAC-3’, GeneDesert (Rev) 5’-TCCAGATGCTGAGAGAAAAACAAC-3’

## Data availability

All data have been deposited with NCBI GEO under accession nos. GEO: GSE306576 (BulkRNA-seq), GEO: GSE306577 (CUT&Tag), GEO: GSE306671 (scRNA-seq) and GEO: GSE306673 (scATAC-seq) are publicly available as of the date of publication.

The code used in this research can be accessed via the following links: https://github.com/ajaykumarsaw/Fasting-primes-small-intestinal-regeneration-after-damage-via-a-microbiome-metabolite-chromatin-axis

## Acknowledgements

The authors thank all members of the Piwnica-Worms and Rai laboratories for constructive criticism throughout the study. We also thank Robert Jenq for his assistance with microbiome work. Funding sources that supported this work include National Cancer Institute of the National Institutes of Health under award number R01CA269495 (to HPW and KR) and Award number RP220567 from the CANCER PREVENTION AND RESEARCH INSTITUTE OF TEXAS (CPRIT) to H.P-W. Experiments performed in this study utilized the MD Anderson Cancer Center microbiome, metabolomics and small animal imaging facility (SAIF) core facility supported by NIH/NCI P30CA016672 grant. We are thankful to MD Anderson Cancer Center advanced technology genomics core (ATGC) facility supported by NIH1S10OD024977-01 and NIH/NCI P30CA016672 grants. We are thankful to MD Anderson Cancer Center the flow cytometry and cellular imaging core facility supported by NIH/NCI 1R50CA243707-01A1 and NIH/NCI P30CA016672 grants. The graphical model was created with BioRender.com (Created in BioRender. Rai, K. (2026) https://BioRender.com/cfwbiov).

## Author Contributions

Conceptualization, P.B, K.R. and H.P.-W.; Methodology, P.B; Software, A.K.Saw., E.A. and K.R; Formal Analysis, A.K.Saw., E.A. and C.C.C.; Investigation, P.B., S.J.J., J.S., A.K.Saw.; E.A. C.C.C., A.K.S., S.S.; Resources, H.P.-W. and K.R.; Writing – Original Draft, P.B. and H.P.W.; Funding Acquisition, K.R. and H.P.W.; Coordination of Study, R.R.J, K.R. and H.P.W. All authors have critically read, edited and approved the final version of the manuscript.

## Declaration of interests

The authors declare no competing interests.

## Notes

### Competing Interest Statement

The authors have declared no competing interest.

