## Supplementary figures for "Fasting primes small intestinal regeneration after damage via a microbiome–metabolite-chromatin axis"

**Supplementary Fig. 1. Fasting significantly alters the gut microbiome.**

Relative abundance of bacteria at genus level determined from 16S rRNA sequencing of ileums isolated on day 0 as shown in Figure 1. Statistical significance was determined by the Mann-Whitney U test.

### Supplementary Fig. 1

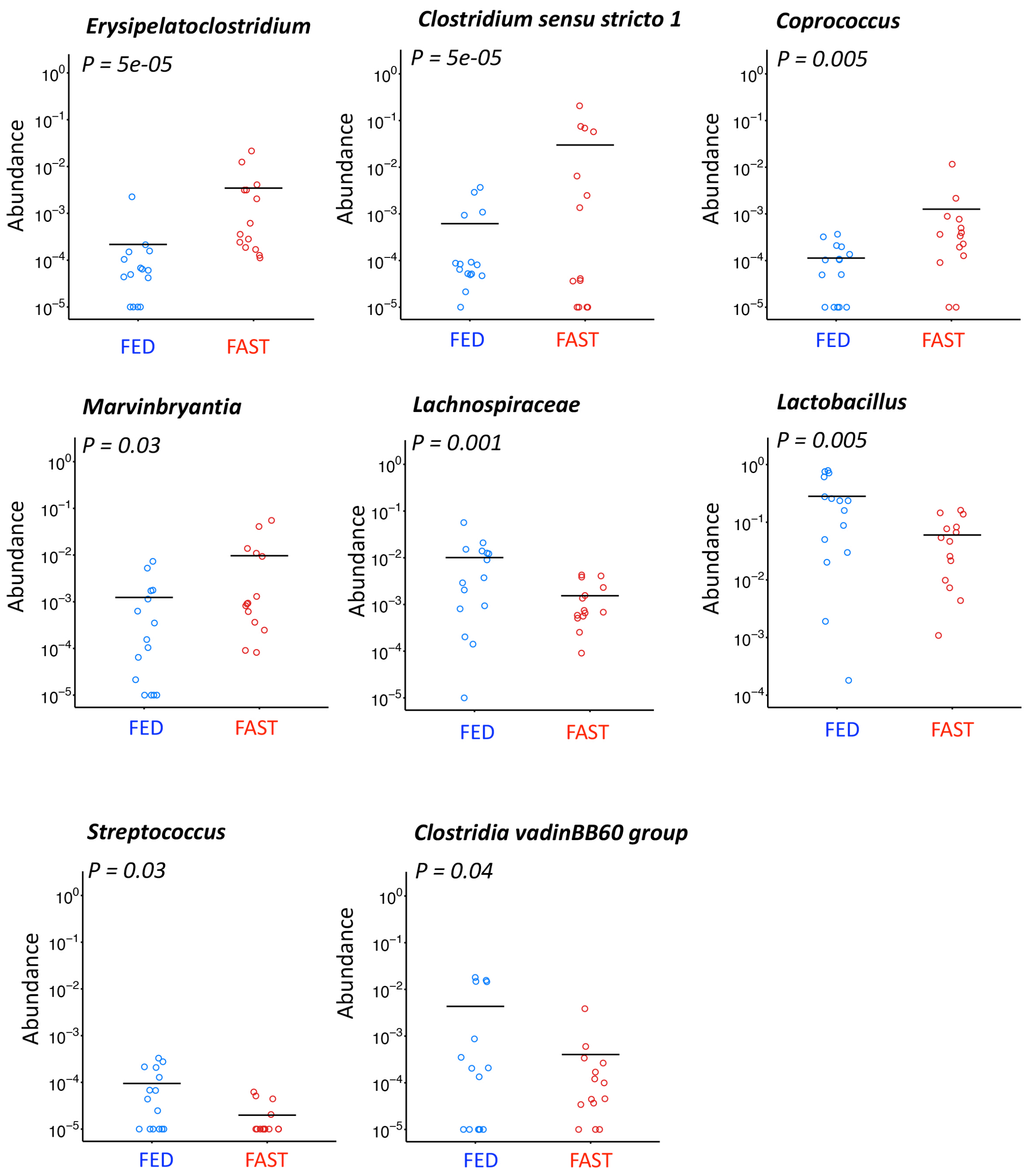

**Supplementary Fig. 2. Broad Spectrum antibiotics significantly alter the gut microbiome.**

a. Diversity of the SI microbiome in ileum samples collected on D-10 (Fig. 2) was quantified using Simpson's reciprocal index. P=1e-05 (Mann-Whitney U test)

b. Microbiome composition determined using principal coordinates analysis based on weighted UniFrac distances in D-10 ileum samples (Fig. 2). Statistical significance was determined by permutational MANOVA testing. *P=0.001*

c. Relative abundances of bacteria at the genus level in ileum samples collected from vehicle- and antibiotic-treated mice on day D-10 (Fig. 2). Each column represents an individual mouse.

**Supplementary Fig. 2**

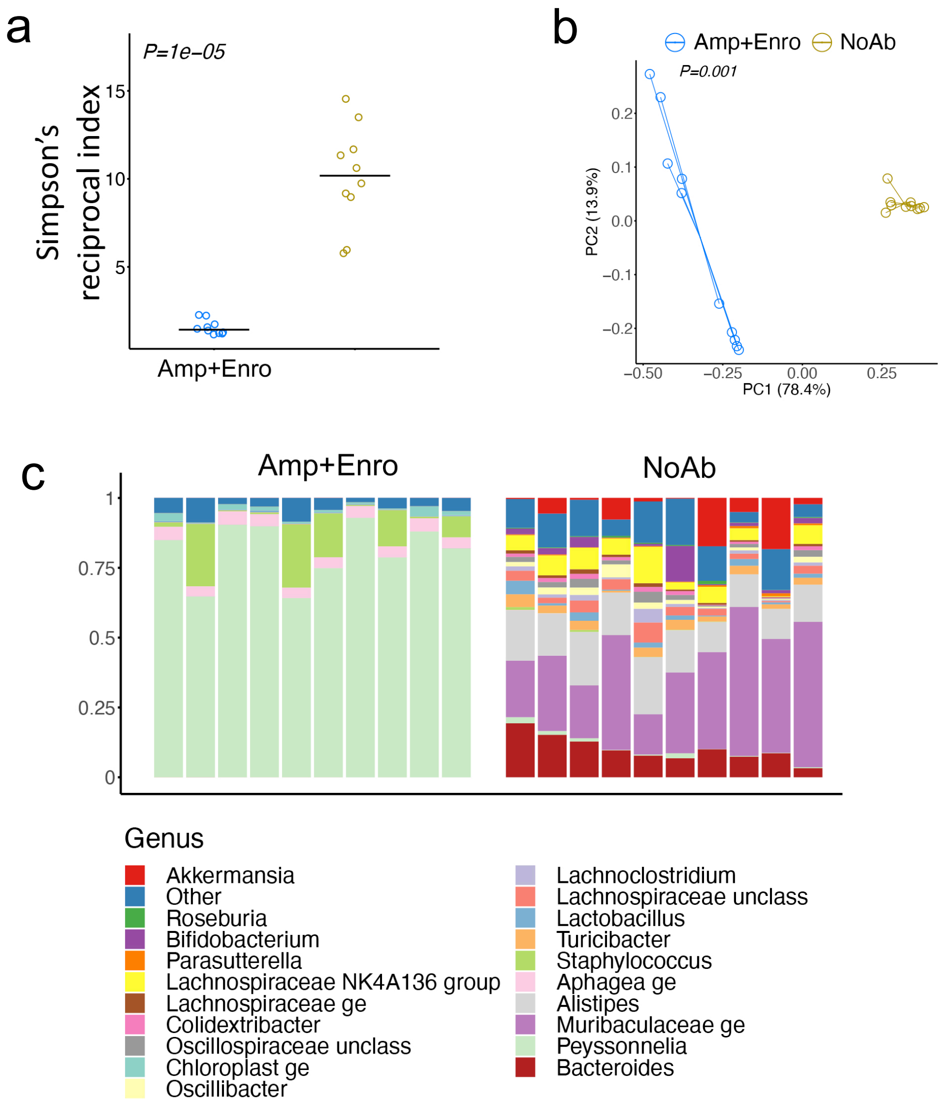

**Supplementary Fig. 3. Tetracycline depletes gut microbiome of *AKK***

a. PCR amplification using primers specific for *AKK* specific genes from day 10 samples isolated from vehicle- and tetracycline-treated mice from experiment shown in Fig. 3. n=5 mice.

b. Raw 16S rRNA read counts for *AKK* in tetracycline- or vehicle-treated mice ileum samples.

c. Relative abundances of bacteria at the genus level in ileum samples collected from tetracycline- or vehicle-treated mice on day -10 (Fig. 3). Each column represents an individual mouse.

**Supplementary Fig. 3**

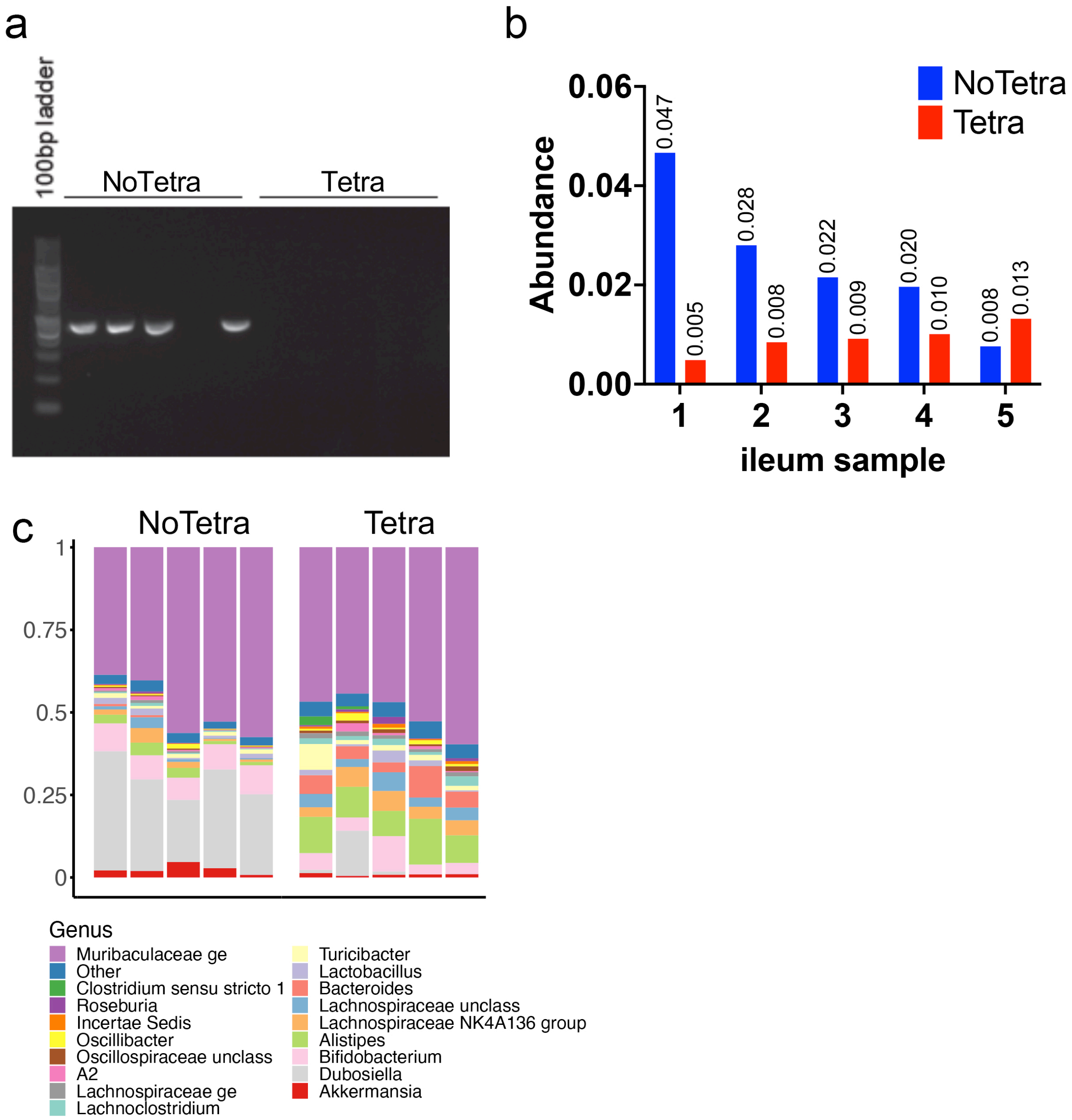

**Supplementary Fig. 4. Single-cell transcriptional profiling on the Fed and Fasted mice small intestinal epithelial cells.**

a. Graphical abstract of experimental methodology. Small intestinal crypt cells were analyzed by single cell RNA-seq.

b. UMAP colored by cluster identity, identifying major intestinal epithelial cell populations.

Supplementary Fig. 4

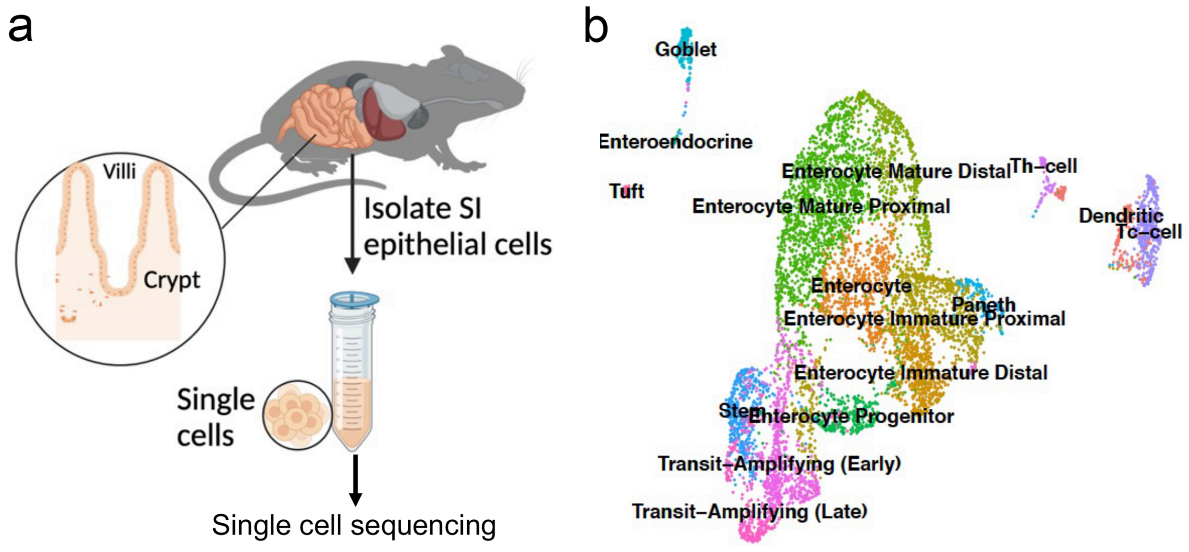

**Supplementary Fig. 5. *AKK* differentially enriches H3K9ac at proximal promoters of apoptotic and proliferative program in small intestine crypts.**

a. Enrichment plot for H3K9ac peaks in Fed (-Tetra), Fed (+Tetra), Fast (-Tetra) and Fast (+Tetra).

b. UpSet plots showing overlap of CUT&Tag seq H3K9ac peaks among Fed (-Tetra), Fed (+Tetra), Fast (-Tetra) and Fast (+Tetra). The CUT&Tag seq peaks unique to Fast (-Tetra) were analyzed separately.

c. Top 10 MSigDB GSEA pathways based on H3K9ac proximal promoter peaks overlapping with publicly available HiChIP and inhouse ChIP seq data in Fast (-Tetra) condition.

d. Genome browser view of CUT&Tag tracks for H3K9ac-enriched regions in SI crypts at p53 pathway-responsive genes (Vdr, Fos and Nupr) under Fast (-Tetra) conditions.

**Supplementary Fig. 5**

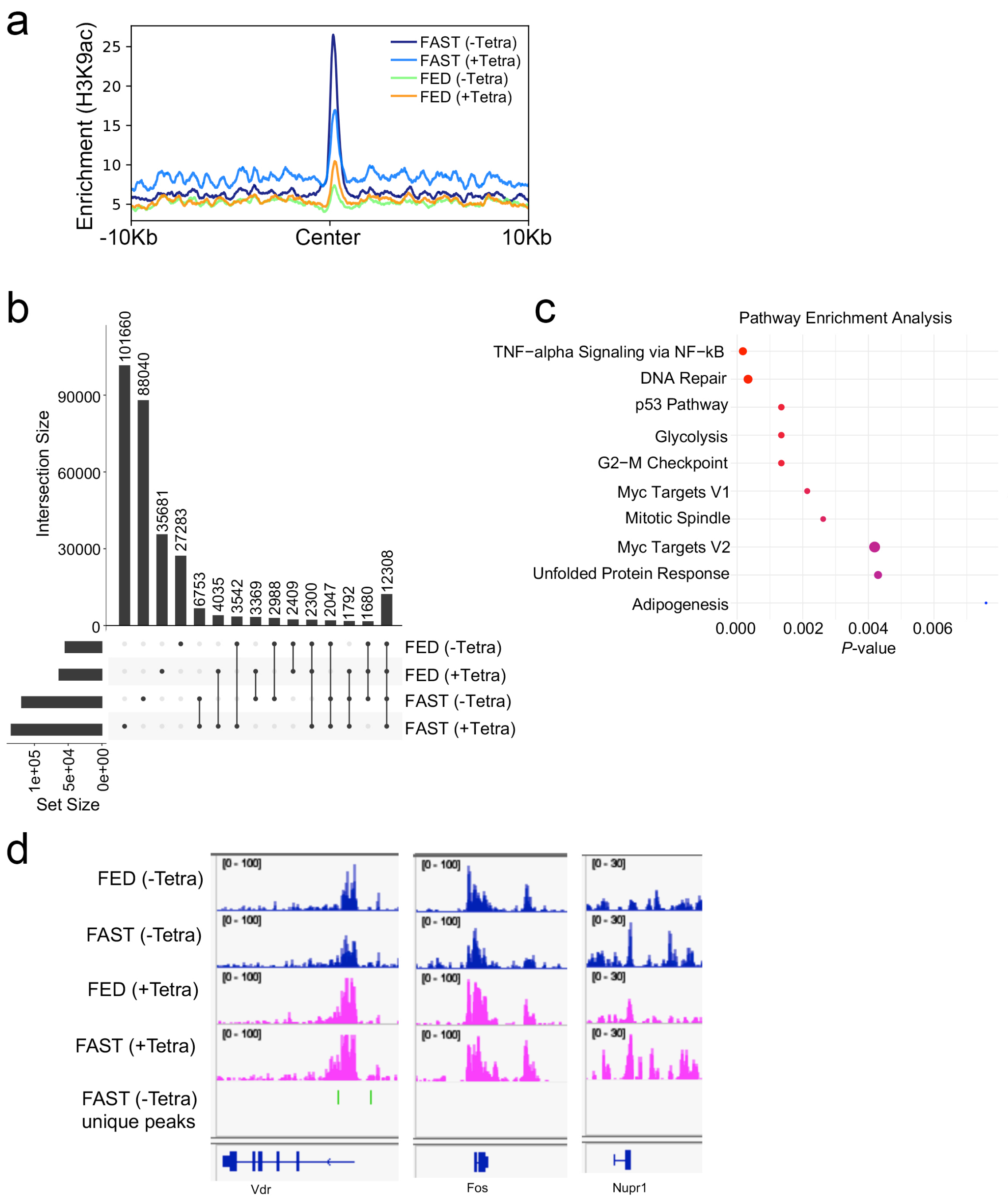

**Supplementary Fig. 6. Fasting remodels the chromatin landscape of intestinal stem and revival cell populations.**

a. UMAP projections of scATAC-seq profiles from fed and fasted mice at 0 h, 24 hpi, 48 hpi, and 72 hpi after irradiation, showing shifts in chromatin accessibility over time.

b. Heat map of scRNA-seq dataset showing gene expression patterns of cluster-enriched markers.

c. IPA canonical pathway enrichment heatmap showing −log10 p-value derived from right-tailed Fisher's exact tests. Pathways with −log10 (p-value) ≥ 1.3 (p ≤ 0.05) were considered significant. Color intensity indicates magnitude of enrichment significance.

d. Genome browser view of scATAC tracks showing representative regions comparing Cluster 3 (blue) and Cluster 6 (magenta).

e. Percentage of Clu high cells (Cluster 3) across all samples.

f. Percentage of stem cells (Lgr5+, Olfm4+), Revival cells (Clu+) and enterocytes (Alpi+) in fed mice and mice fasted for 24h at 0 h post-irradiation assessed nby single cell RNA sequencing.

g. Integration of CUT&Tag H3K9ac data with RNA-seq for the activated genes in Fed (-Tetra), Fed (+Tetra), Fast (-Tetra) and Fast (+Tetra) samples. All genes displayed are differentially expressed (p ≤ 0.05, log2FC ≥ 2 or ≤ –2). n = 2 mice per group.

### Supplementary Fig. 6

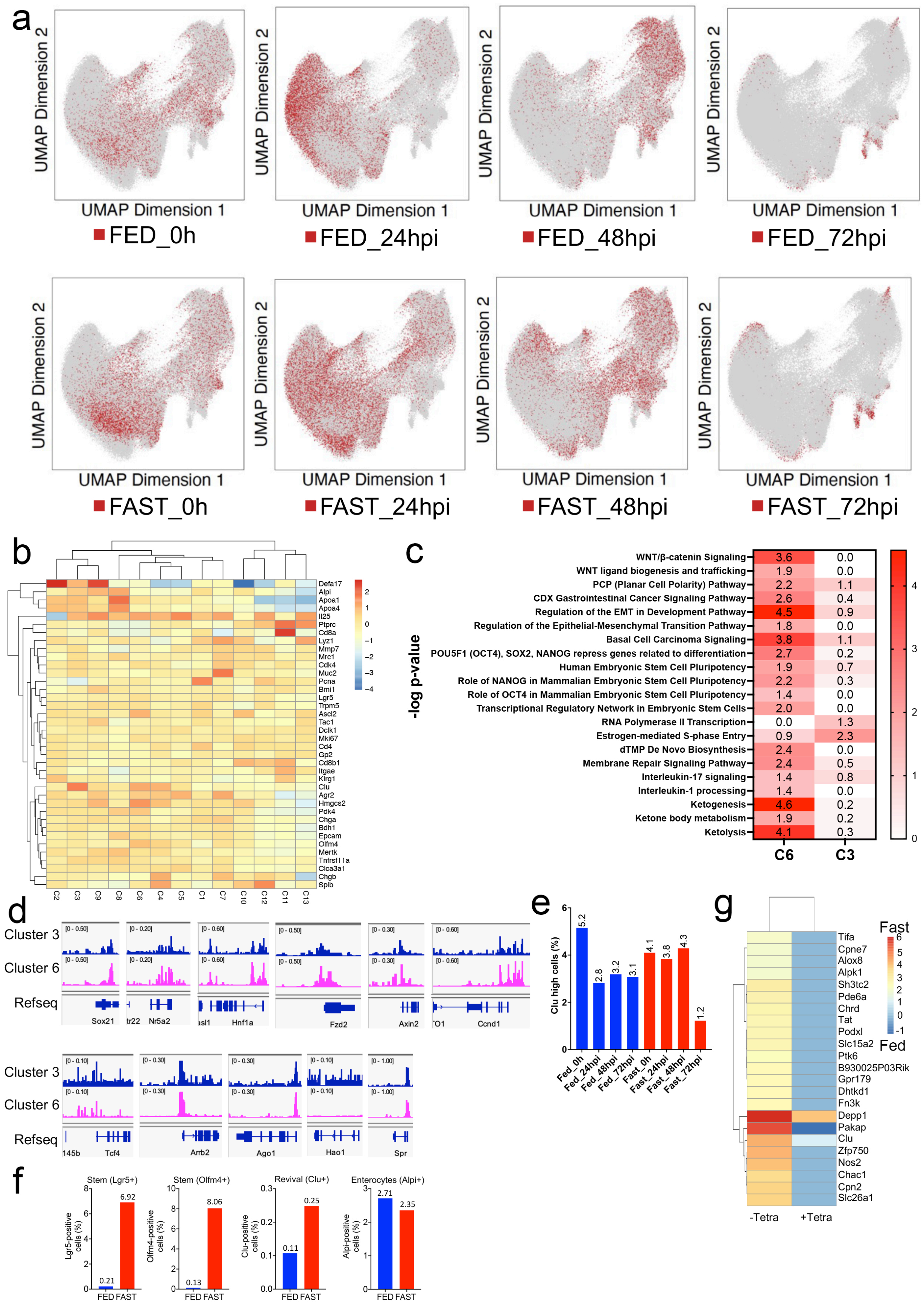

**Supplementary Fig. 7. scATAC-seq analysis of Clu-associated cell states.**

scATAC-seq–based chromatin trajectory analysis shown on UMAPs for fed and fasted mice at 0 h, 24 hpi and 48 hpi. Trajectory C6 (Clu+Olfm4+) -> C3 (Clu high) -> C1 (TA cell) -> C8 (Enterocyte).

**Supplementary Fig. 7**

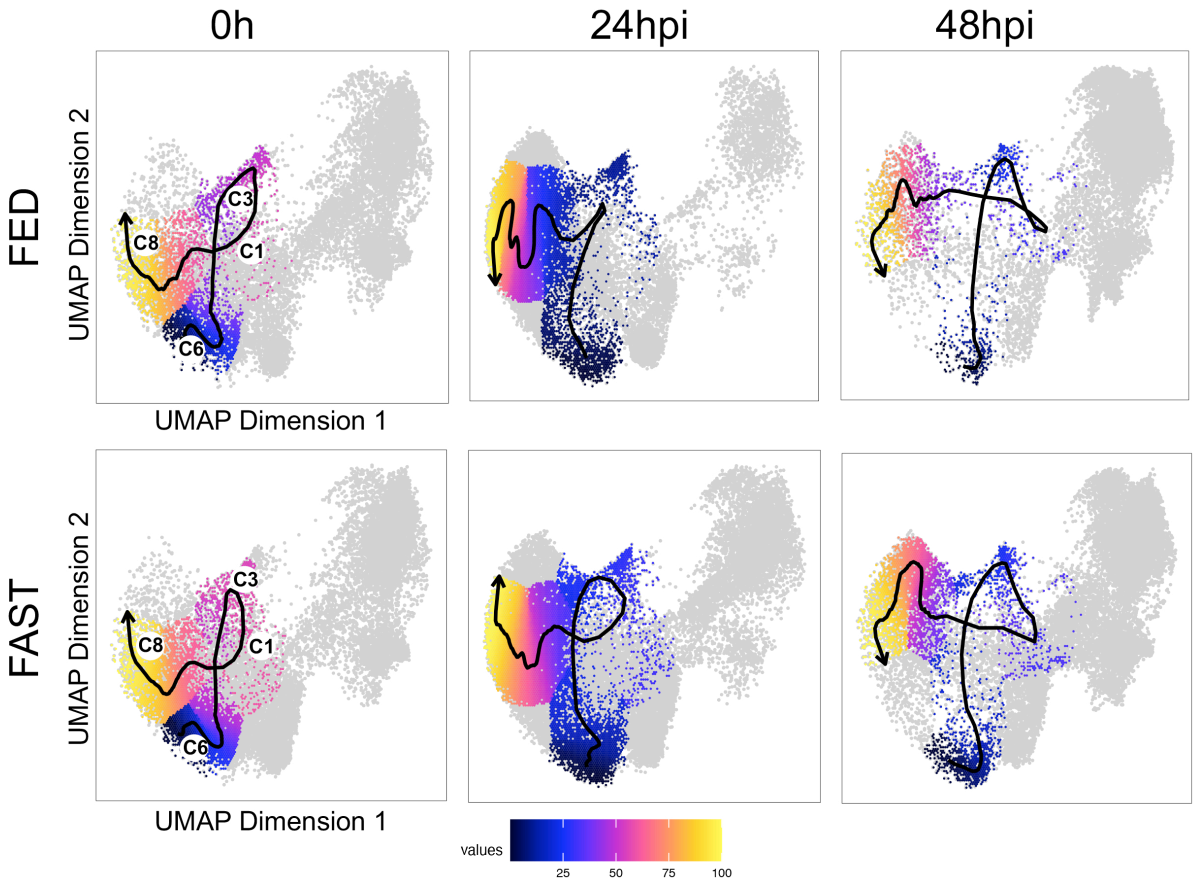

**Supplementary Fig. 8. scATAC-seq reveals time-resolved chromatin dynamics in fed and fasted intestinal epithelium.**

a. Bar graph showing the number of shared accessible regions between scATAC-seq and H3K27ac CUT&Tag fed datasets. “Common” refers to overlapping enhancer-like peaks.

b. Correlation between *Clu* chromatin accessibility and transcription factor motif accessibility across single cells.

**Supplementary Fig. 8**

a

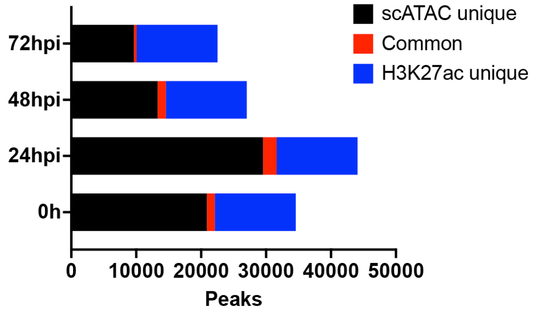


b

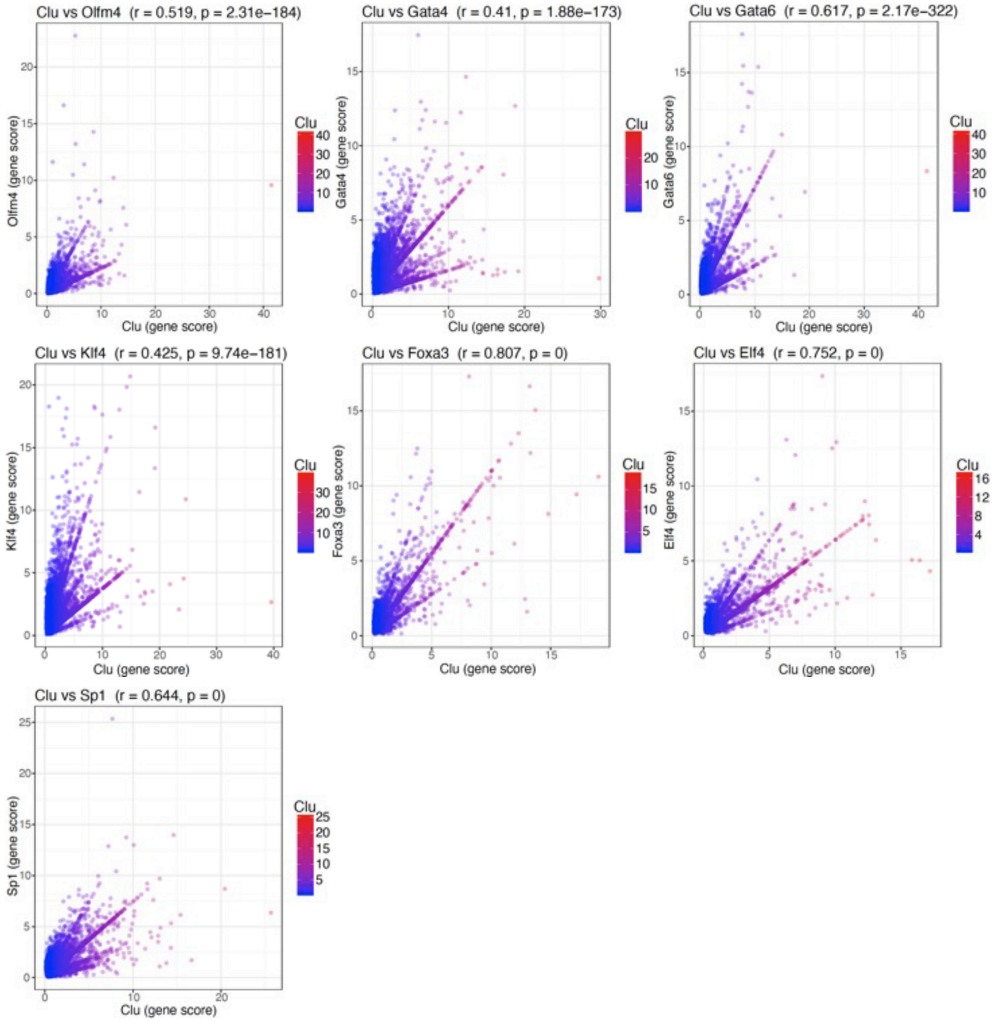
